# *Dmrt2* Regulates Sex-Biased Neuronal Development In The Cingulate Cortex

**DOI:** 10.1101/2025.01.08.631875

**Authors:** Ana Bermejo-Santos, Miguel Rubio-García, Rodrigo Torrillas-de la Cal, Rafael Casado-Navarro, Esther Serrano-Saiz

## Abstract

Sexual differences are prevalent in the brain. DMRT transcription factors have been postulated as important determinants of sex differences. Previous research focused on the DMA subfamily in the brain. Here, we reveal an unprecedented role for *Dmrt2* in regulating the proliferation and development of cortical neurons in mice. *Dmrt2* is expressed in deep-layer neurons of the cingulate cortex (CgCx) throughout development. Its downregulation in the CgCx primordium results in premature cell cycle exit of embryonic progenitors and subsequent reduction in cortical plate cellular density at later developmental stages. *Dmrt2* expression is higher in male embryos during early development, potentially explaining their increased vulnerability to *Dmrt2* depletion. As development progresses, *Dmrt2* expression persists in deep-layer neurons, controlling processes like migration, axonal targeting, and neuronal-specific gene expression. This study broadens our understanding of *Dmrt2* gene function in the brain and provides insights into the molecular basis of sexual differences in neurodevelopmental processes.

## INTRODUCTION

Sex differences are prominent in nearly every neuropsychiatric and neurodevelopmental disorder, influencing the age of onset, prevalence, symptomatology, diagnosis, and therapeutic response^1,2^. The genetic factors involved in the sexual differentiation of the brain could underlie the sexual bias and etiology of psychiatric disorders. The *<u>d</u>oublesex and <u>m</u>ab-3-related transcription factor* (*Dmrts*) gene family stands out for its evolutionary conservation as key regulators of sex determination and differentiation^3–5^. The family is defined by the region that mediates DNA binding, the DM domain, which is in all *Dmrts*^6^. A DMRT subgroup has an additional domain, the DMA domain of unknown function. *Dmrts* have been extensively studied in the sexual differentiation of invertebrate nervous systems (NS) and have been implicated in the control of cell death, terminal identity differentiation, and synaptic connectivity to, ultimately, generate sex-specific circuit configurations^7,8^. However, we currently lack a full understanding of the role of *Dmrt* genes in sexual differentiation within mammalian NS.

In mammals, a role in the NS has only been addressed for the subfamily of DMA-*Dmrts* (including *Dmrt3*, *Dmrt4,* and *Dmrt5*) (reviewed in^9^). Work in vertebrate embryos (mice and *Xenopus Laevis*) shows that, depending on the cellular context, DMA-*Dmrts* are involved in maintaining the proliferative capacity of neural progenitor cells in the mouse medial telencephalon^10,11^ or neuronal differentiation processes^12–14^.

Besides the DMA subfamily, other family members, including *Dmrt2*, are also expressed in the mouse brain, but no neuronal function has been attributed to them yet. The cingulate cortex (CgCx) primordium is one of the earliest regions where *Dmrt2* is detected. *Dmrt2* has been previously identified as expressed in postmitotic cingulate corticothalamic projection neurons (CThPNs), where it is proposed to exert late developmental functions, such as synaptic targeting and maturation^15^. *Dmrt2* functions have been exclusively described outside the NS, where it plays important roles in cell type differentiation, particularly in somite development. Here, *Dmrt2* regulates dermomyotomal and myotomal transcription factors, including PAX3 and MYF5, required to initiate skeletal muscle formation^16,17^. In addition, *Dmrt2* is a key player in chondrocyte differentiation during bone formation. *Dmrt2* acts as a *Sox9*-inducible gene that promotes the transition from a proliferative to a more differentiated hypertrophic stage^18^.

Although little is known about the function of *DMRT2* in the human brain, several studies have shown that deletion or duplication of the short arm of chromosome 9, where *DMRT1*, *DMRT2*, and *DMRT3* are located, is associated with several mental disorders or intellectual disabilities. These include developmental delay and craniofacial anomalies, autism spectrum disorder^19,20^, and obsessive-compulsive disorder^21,22^. More specifically, *DMRT2* has been postulated as a risk factor for obsessive-compulsive disorder^23^. While no studies have specifically identified the neuronal function of DMRT2, one report conducted in human neuroblastoma cells shows that DMRT2 regulates both the proliferation rate and the progression of tumor growth^24^. However, the mechanisms by which *DMRT2* may induce or influence these processes *in vivo* have not yet been described.

In this study, we have analyzed the role of *Dmrt2* in the development of the mouse cingulate cortex, a region critical for emotional processing, executive function, and social behavior. By comparing male and female mouse embryos, we studied a novel link between *Dmrt2* function and the sexual differentiation of the mammalian NS. First, we thoroughly examined *Dmrt2* expression in the mouse CgCx at embryonic and postnatal time points in males and females. We asked whether there are sexual and developmental differences between *Dmrt2* splicing variants predicted by the NCBI genome browser (NCBI Gene ID: 226049). Then, we performed a functional analysis by downregulating, with interference RNA (iRNA), or overexpressing *Dmrt2* through *in utero* electroporations (IUE). We compared transcriptional profiles of male and female embryos and the general progression of proliferation, migration, and differentiation of cingulate cortical neurons. Our results provide valuable insights into developmental neurobiology and sex-based neuroscience by exploring the intersection of *Dmrt2* function, sex-specific development, and cortical formation.

## RESULTS

### *Dmrt2* is expressed in mouse deep-layer cingulate cortical neurons

We analyzed the expression of *Dmrt2* by *in situ* hybridization (ISH) in male and female embryos (E) at E12.5, E13.5, E14.5, and E18.5, and postnatal (P) 7 (**Figure 1A-1D, Suppl. Figure 1A-1D**) and adults P56 animals (**Suppl. Figure 1D**). We used a previously published riboprobe encompassing the coding RNA sequence^17^ (**Figure 1E**).

**Figure 1.**
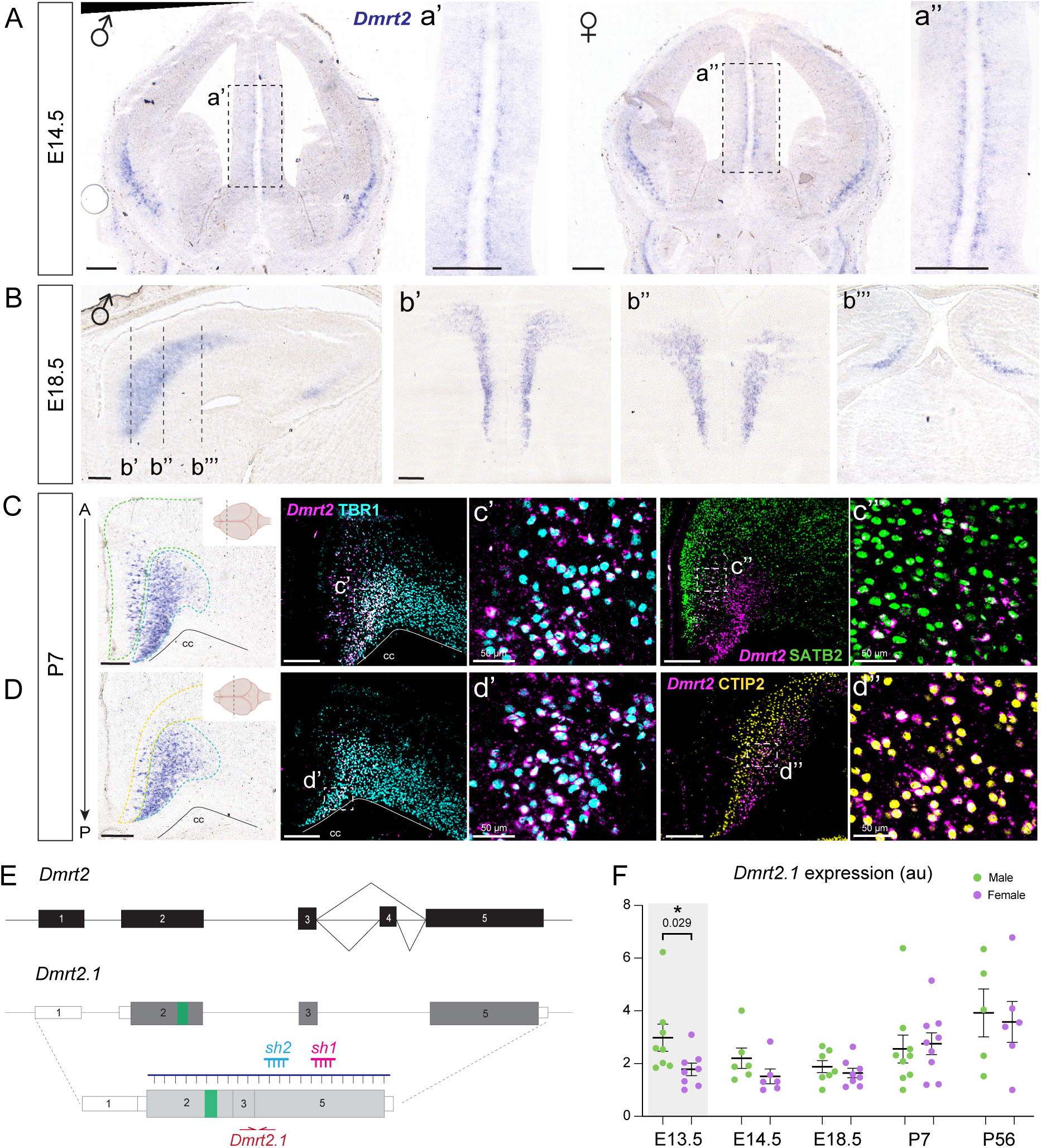
*Dmrt2* is expressed in the deep-layer neurons of the cingulate cortex at higher levels in male mice during early cortical development. In all panels, *Dmrt2* was detected by ISH **A)** *Dmrt2* in the mantle zone of male and female embryos at E14.5. (a’, a’’). Insets show a higher magnification of the cingulate primordium. **B)** By E18.5, ***Dmrt2* occupies the entire cingulate cortex in its rostrocaudal axis.** Only male images are depicted. Each coronal section in b’-b’’’ corresponds to the sagittal level indicated with a dashed line in the left panel. **C-D) *Dmrt2* expression in postnatal cingulate cortices at P7.** In the anterior cingulate cortex, *Dmrt2* overlaps with TBR1 (c’, d’) and SATB2 in more superficial layers (c’’). Additionally, in the mid-cingulate, *Dmrt2* colocalizes with CTIP2 (d’’). cc: corpus callosum. Scale bar: 250 μm, except in the indicated cases. **E) *Dmrt2* locus schematic.** Solid black and grey boxes represent exons; white boxes represent untranslated regions (UTR); the green box within exon 2 indicates the location of the DM domain; purple line, *Dmrt2* riboprobe used for ISH detection; blue and pink line, interference RNAs used in this study to target *Dmrt2*. **F) *Dmrt2.1* splicing variant is higher in males in early development.** qPCR analyzed cingulate cortices of male and female embryos with the primers depicted in E). Red arrows at several time points throughout development. The graph represents mean±SEM in arbitrary units (au). Each dot corresponds to one individual embryo. Two-tailed t-test: * p-value < 0.05 (n ≥ 5).

*Dmrt2* ISH at E12.5 shows a strong signal in the respiratory epithelium (RE). By E13.5 a few positive cells can be seen in the ventral cortex (**Suppl. Figure 1B**). By E14.5, *Dmrt2* is robustly detected in the cingulate cortex (CgCx) primordia, and it is still maintained in the ventral cortex in both sexes (**Figure 1A**). *Dmrt2* occupies the CgCx in its entire rostrocaudal axis, including the retrosplenial cortex (RSP) and excluding the somatosensory cortex. Additionally, it can be detected in distinct areas of the neocortex, such as the prelimbic (PLC) and the infralimbic (Ila) areas (**Figure 1B**; **Suppl. Figure 1C**). We look at P7, when migration is complete, to identify *Dmrt2* CgCx neurons through double immunofluorescence and ISH. We use TBR1 as a general marker of deeper layer VI^25^ and CThPN identity^26^ and SATB2 as a marker of upper callosal projection neurons (CPN) (layer II to VI, although expression in layer VI is dimmer)^27^. *Dmrt2*-labeling is strong in layer VI-TBR1+ cells across the rostrocaudal length of the CgCx (**Figure 1C, c’, d’**). An additional superficial cluster is detected in the anterior CgCx, where *Dmrt2* colocalizes with SATB2 in layer V CPN (**Figure 1C, c’’**). At postnatal stages, a population of *Dmrt2+* cells is clearly found in layer V, defined by CTIP2^26^ (**Figure 1C, d’’**). In summary, *Dmrt2* is primarily expressed in cingulate, deeper CThPNs from mid-gestation embryos until postnatal animals and in a few scattered CPNs.

### *Dmrt2.1* is expressed at higher levels in males during early cortical development

The NCBI genome browser shows that the *Dmrt2* locus has three alternative transcript variants (Gene ID: 226049). According to predictions, *Dmrt2.2* and *Dmrt2.3* retain exon 4, which contains an alternative stop codon. *Dmrt2.2* is predicted to produce a shorter protein containing a longer 3’UTR than *Dmrt2.1* or *Dmrt2.3*. *Dmrt2.1* and *Dmrt2.2* sequences include the DM domain (located in exon 2, indicated in green in **Suppl. Figure 2A**). On the other hand, *Dmrt2.3* mRNA will be transcribed from a cryptic transcription start site in exon 2, but its translation will commence from an alternative start site in exon 5. This protein product will lack the DM domain and share the 3’UTR with *Dmrt2.1* (**Suppl. Figure 2A**). Whether or not these variants are expressed in the mouse NS has never been addressed.

To test whether there is a sex-specific effect on *Dmrt2* expression levels, we performed real-time quantitative PCR (qPCR) at E13.5, E14.5, E18.5, P7, and P56 stages. We included samples from antero-dorsal cortices containing the CgCx from male and female (we excluded the hippocampus from E18.5 and postnatal brains; see **Suppl. Figure 1B,1C** for the dissection area). We designed primers that detect *Dmrt2.1 and Dmrt2.2* variants (“*Dmrt2-Ex1*”), *Dmrt2.1* specifically (“*Dmrt2.1”)*, or *Dmrt2.2* and *Dmrt2.3* (“*Dmrt2.2/2.3*”) (**Figure 1E**; **Suppl. Figure 2A**). At E13.5, *Dmrt2.1* is significantly more expressed in male embryos than in females (1.67 times more); however, at later developmental stages, we no longer detected any sexual difference between the two sexes (**Figure 1F; Suppl. Figure 2C**). Only *Dmrt2.1* variant shows a differential expression between male and female animals, while *Dmrt2.2/2.3* are similarly expressed in the two sexes at all developmental time points (**Suppl. Figure 2D**). Thus, we focus our study on *Dmrt2.1* from now on.

### *Dmrt2* downregulation triggers CgCx precursors’ premature exit from cell cycle more substantially in male embryos

To investigate the function of *Dmrt2* in the nervous system, we conducted a series of experiments to manipulate *Dmrt2* expression specifically in the CgCx through *in utero* electroporation (IUE). We electroporated a plasmid coding for a short hairpin (sh) RNA (designated sh*Dmrt2*; sh1 in **Figure 1E, Suppl. Figure 2A**), and compared it to mock conditions (see **Experimental Procedures** for details). We confirmed that the sh*Dmrt2* induces *Dmrt2* downregulation after 5 days of electroporation, at E18.5, by measuring *Dmrt2* mRNA levels of the ipsilateral versus the contralateral cingulate cortex (**Suppl. Figure 2F**).

We electroporated the sh*Dmrt2* at E13.5, when deeper-layer neurons (DLN) are born^28^, to investigate early effects on neurogenesis and proliferation. Two days after electroporation, at E15.5, we observed a striking reduction in GFP(+) cells in sh*Dmrt2*-electroporated cingulate primordia (**Figure 2A-2D**). We quantified the **total cellular density** of GFP(+) cells across the cingulate primordium (number of GFP(+) cells/μm^2^): GFP(+) are reduced in sh*Dmrt2* males from 2.63±0.12 vs. 1.35±0.29 10^-3^ cells/μm^2^; however, the effect is not significant in female embryos, although it follows a similar trend (2.30±0.35 vs 1.85±0.36 10^-3^ cells/μm^2^) (**Figure 2E**).

**Figure 2.**
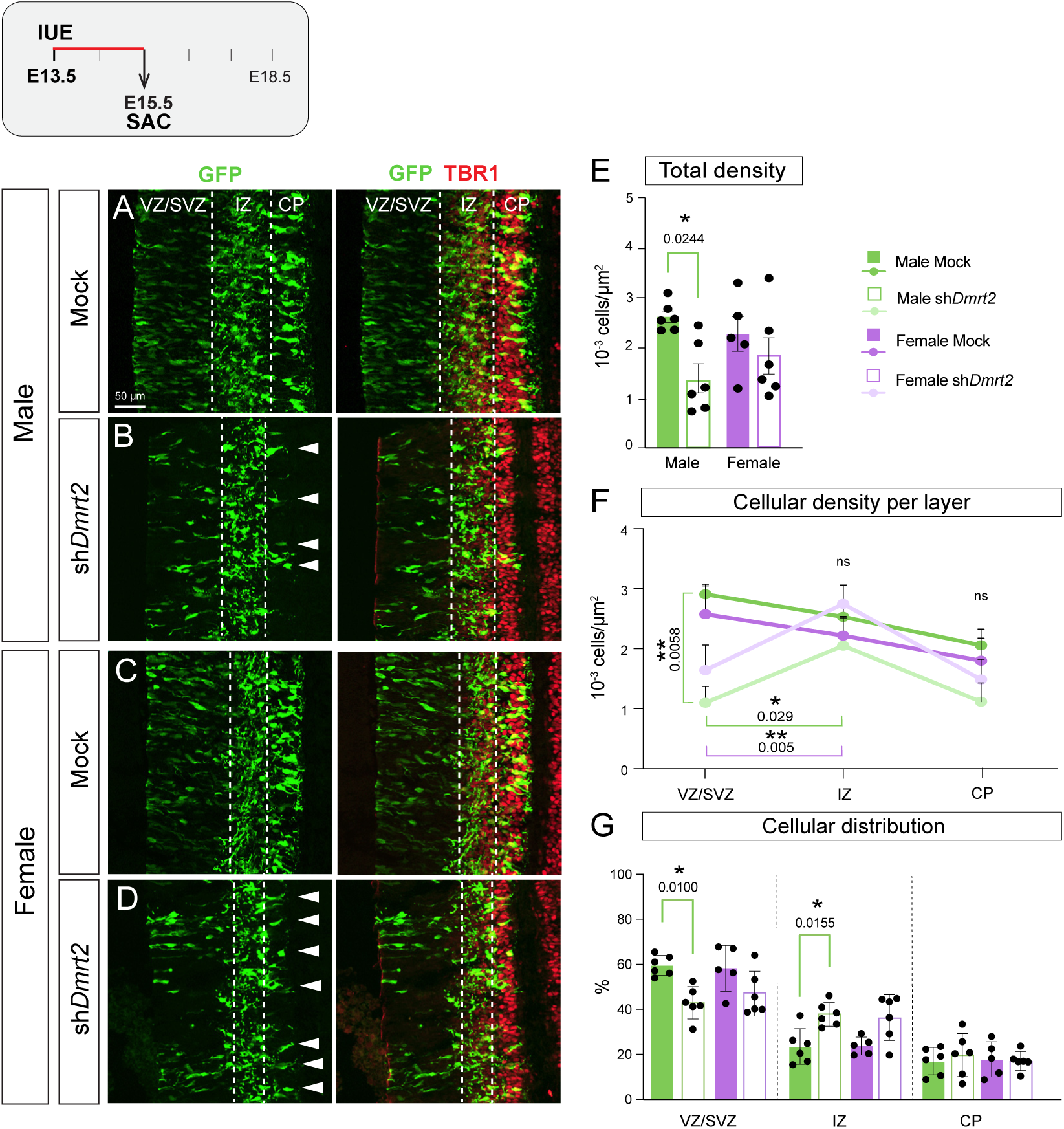
*Dmrt2* downregulation affects the proliferation of CgCx progenitors more strongly in male embryos. **A-D)** E15.5 CgCx primordium of electroporated embryos at E13.5. GFP(+) cells populate the three layers VZ/SVZ, IZ and CP (TBR1+). **A) Male mock, B) Male sh*Dmrt2*, C) Female mock, D) Female sh*Dmrt2***. White arrowheads in B and D point to the somas of the few GFP(+) cells that reached the CP and indicate a stronger effect in males quantified in E-G. **E) Total GFP cellular density within the three layers.** Tukey’s multiple comparison test, ordinary one-way ANOVA (n ≥ 5). **F) Cellular density in each layer in the four conditions.** Tukey’s multiple comparison test, ordinary one-way ANOVA (n = 6) was applied to compare values within layers. Regression analysis was conducted to compare the cellular density progression between VZ and IZ layers, and VZ and CP layers. **G) Quantifications of GFP(+) cellular distribution (%)**. Dunn’s multiple comparison test, Kruskal-Wallis test (n ≥ 5). For all graphs, each dot corresponds to one individual embryo; results are expressed as mean ± SEM, and significances are as follows: * p-value < 0.05 and ** p-value < 0.01. ns: not significant.

The substantial reduction in the total GFP(+) cells could result from reduced proliferation, increased neurogenesis, or cell death (which we discarded because of negative cleaved-Caspase3 signal and TUNEL. Data not shown). At E15.5, GFP(+)-DLNs had time to migrate to the cortical plate (CP), and we can analyze the distribution across the CgCx primordium layers: the ventricular zone (VZ), where apical progenitors reside (Ki67+); the subventricular zone (SVZ) marked by TBR2, the intermediate zone (IZ), where cells are migrating and axons growing, and the cortical plate (CP), where mature neurons migrate, marked by dim TBR1 and high TBR1, respectively^29^ (**Suppl. Figure 3A**). Quantitative analysis of **cellular densities** across cortical layers revealed altered developmental trajectories in sh*Dmrt2* embryos (**Table 1)**. In mock controls, GFP(+) cells predominantly localize to the VZ. As cells differentiate, they progressively populate the IZ and CP. At E15.5, the CP is the least populated layer (**Figures 2F, 2G)**. In contrast, sh*Dmrt2* embryos exhibited an altered developmental dynamic, characterized by the aberrant increase of GFP(+) cells in the IZ at the expense of VZ and CP populations (**Figure 2F**). In sh*Dmrt2* embryos, GFP(+) cell densities are significantly reduced in the VZs compared to mock controls. Conversely, the IZ exhibits increased cellular accumulation in sh*Dmrt2* embryos relative to their VZs, whereas mock conditions show the inverse pattern (IZ less densely populated than VZ; **Figure 2F**). Statistical regression analysis of cellular density progression revealed a significant divergence between sh*Dmrt2* and mock embryos in the VZ-to-IZ transition, with this effect being consistent across both sexes. The VZ-to-CP transition showed no statistically significant differences between experimental groups. These results suggest that progenitors in the VZ are prematurely exiting the cell cycle.

Although both sexes presented a similar altered progression, there was a higher reduction in the total cellular densities in male sh*Dmrt2* compared to male mock embryos. The quantification of the **cellular distribution** of GFP(+) cells in each layer (**Figure 2F**; **Table 1**) revealed a significant reduction in GFP(+) in the VZ of sh*Dmrt2* relative to mock in male embryos, while the reduction is not significant in females. Concomitantly, there is an increase in the proportion of IZ-GFP(+) cells in sh*Dmrt2* males vs. mock males, but not in females. Lastly, CP-GFP(+) cells show a slight, non-significant increase in their proportions in sh*Dmrt2* relative to mock males. In sum, *Dmrt2* downregulation affected male embryos more strongly, although similar processes seem to operate in both sexes.

At E15.5, *Dmrt2* is robustly detected in the CP by ISH, and we could not detect *Dmrt2* transcripts in the VZ/SVZ or the IZ (**Suppl. Figure 3B**). However, we confirmed that the phenotype observed in this study is specific to *Dmrt2* by electroporating a distinct sh*RNA* (sh2, **Figure 1E**) and obtaining similar results (**Suppl. Figure 3C**). Moreover, we took the opposite approach and overexpressed the canonical *Dmrt2.1* through IUE at E13.5 (oe*Dmrt2.1*). After two days from electroporation, at E15.5, the GFP(+) cellular distribution shows the opposite phenotype to the sh*Dmrt2* conditions: *Dmrt2.1* overexpression caused a substantial decrease in CP-GFP(+) cells at the expense of an increase in the VZ-GFP(+) and IZ-GFP(+) cells (**Table 1**. **Suppl. Figure 3D-3H**), indicating that *Dmrt2.1* overexpression caused an increase of progenitors proliferation.

To confirm that *Dmrt2* downregulation triggers the premature exit from the cell cycle, we electroporated embryos at E13.5, but analyzed them at E14.5 (24 hours after IUE) (**Figure 3**). Two hours before collecting the embryos, we injected EdU into the pregnant females to assess cellular proliferation in sh*Dmrt2* and mock electroporated CgCx embryonic progenitors. At E14.5, we found no differences in total GFP(+) cellular densities between sh*Dmrt2* and mock embryos, in any sex (**Figure 3B**). Most cells populate the VZ and SVZ layers, and only a few reach the CP (marked by TBR1). When we analyzed the number of EdU+/GFP+ in each layer, we found a significant reduction in the percentage of EdU+/GFP+ in the SVZ of sh*Dmrt2* brains compared to mock, suggesting that there is a reduction in the number of progenitors in sh*Dmrt2* conditions. We could not detect any differences in cleaved-Caspase3 staining in the CgCx of sh*Dmrt2* or mock brains at this time point. Together, the decrease in EdU(+)/GFP(+) in the SVZ and the increase in the IZ at E15.5 (**Figure 2**) indicate that the downregulation of *Dmrt2* causes a premature exit from the proliferative state.

**Figure 3.**
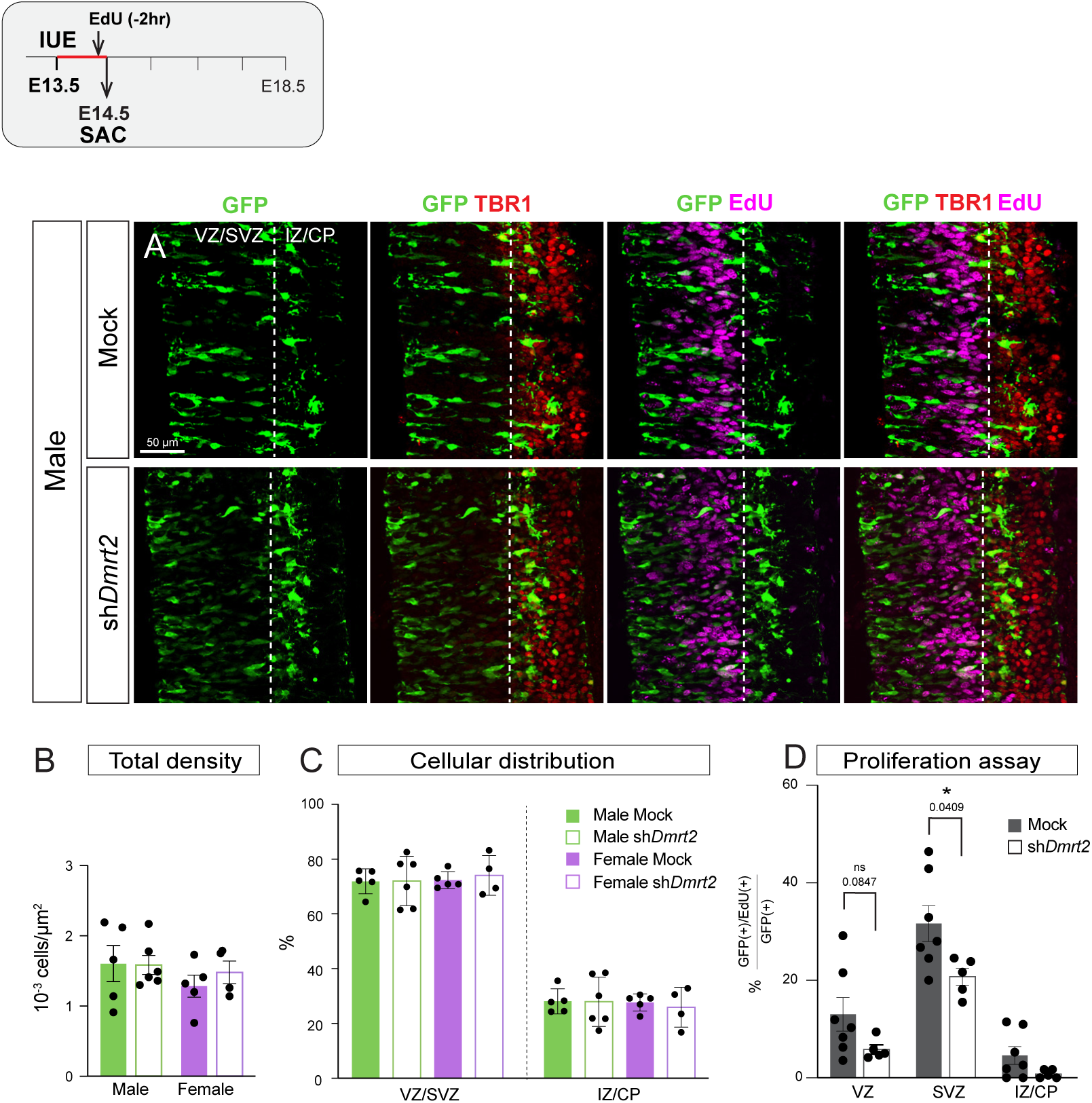
*Dmrt2* downregulation reduces the percentage of progenitors in both sexes. **A)** E14.5 CgCx primordium of electroporated embryos at E13.5. GFP(+) cells are in the VZ/SVZ and incipient IZ/CP (marked by TBR1+). Only one sex is shown. **B) Total GFP cellular density.** Tukey’s multiple comparison test, ordinary one-way ANOVA (n = 5). **C) Quantifications of GFP(+) cellular distribution (%)**. Dunn’s multiple comparison test, Kruskal-Wallis test (n = 5). **D) Proliferation assay.** The percentage of GFP+ and EdU+ cells is quantified for mock and sh*Dmrt2* embryos. Both sexes are pooled. For all graphs, each dot corresponds to one individual embryo; results are expressed as mean ± SEM, and significances are as follows: * p-value < 0.05 and ** p-value < 0.01.

Altogether, *Dmrt2* downregulation is prominently affecting the VZ layer, depleting the neural precursor pool, followed by an increase in cells at the IZ layer. Moreover, *Dmrt2* downregulation affects male embryos more strongly than female embryos. The strong reduction of GFP(+) cells in the VZ suggests that *Dmrt2* is required to maintain progenitor cells in a proliferative state. When it is downregulated, progenitor cells exit the cell cycle prematurely. However, after they exit the cell cycle, GFP(+) cells tend to accumulate in the IZ and do not adequately reach the CP. *Dmrt2.1* overexpression triggers the opposite effect and keeps progenitors in a proliferative state.

### *Dmrt2* is required in post-mitotic neurons for migration and axon guidance

The robust expression of *Dmrt2* in post-mitotic neurons at late developmental stages prompted us to investigate phenotypes in late E18.5 embryos (**Figure 1**, **Suppl. Figure 1**), when the migration of deeper neurons should be mostly complete and axonal extension is taking place. We then compared these results with mock electroporated brains (see **Experimental Procedures** for details).

In mock electroporated animals at E18.5, GFP(+) cells populate homogeneously all layers of the CgCx and extend projections ipsilaterally towards the thalamus dorsally to the internal capsule, while a subset of axons crosses the midline through the corpus callosum (cc) (**Figure 4A, 4C; Suppl. Figure 4A, 4C**). In sh*Dmrt2* embryos, GFP(+) cells are also distributed along the CgCx, although they show several aberrant phenotypes (**Figure 4B, 4D**; **Suppl. Figure 4B, 4D; Table 2; Suppl. Table 1, 2**). **1)** as an indication of the total amount of electroporated cells, we quantified the GFP(+) fluorescence intensity (integrated density (IntDen)) of the whole electroporated area of the cingulate CP (excluding the IZ and the VZ/SVZ) (red dashed line in **Figure 4A-4D**). We found a 33% reduction in males (15% non-significant reduction of fluorescence intensity in females), (**Figure 4E**), which was expected if we take into account the exhaustion of the progenitor pool (**Figures 2, 3**). **2)** We analyzed the GFP(+) cell distribution across the cortical layers in the four conditions. We selected a representative region of interest to avoid “gfp holes” and the medial CgCx, where the cytoarchitecture of the CgCx is less stratified, and divided it into 10 bins (red solid boxes in **Figures 4a’-d’**). We measured the fluorescence intensity (IntDen) within each bin. Bins 10-9 correspond to deeper layers just above the cc, while bins 1-3 comprise the most external layers containing the newly generated neurons that had migrated the most. TBR1 highly expressing neurons (deeper layer VI) correspond roughly to intermediate bins 8-6, and layer V to bins 5-4. Most GFP(+) fluorescence accumulates in mock embryos in intermediate layers. However, in sh*Dmrt2* male embryos, bins closer to the cc show a significant increase in fluorescence at the expense of intermediate bins (**Figure 4F**). This phenotype is observed in both sexes but is more exacerbated in females. In sh*Dmrt2* brains, we found GFP(+) cells stacked in the cc along the whole anterio-posterior axis of the CgCx (white arrowheads in **Figures 4b’, 4d’; Suppl. Figure 4B, 4D**). By quantifying the total number of cells “misplaced” at the cc, we found that the effect is significantly higher in females (**Figure 4G**; **Table 2; Suppl. Table 1, 2**). In both sexes, those cells colocalize with TBR1 and CTIP2, but only some with SATB2 (**Suppl. Figure 4E**), suggesting migration defects. These cells are never found in mock animals. **3)** We systematically observed that the VZ/SVZ is almost completely depleted of GFP(+) cells in the sh*Dmrt2* brains (**Figure 4a’’-d’’; Table 2; Suppl. Table 1, 2)**. The lack of GFP(+) cells in the VZ/SVZ progenitor layer confirms the depletion in the progenitor pool in the sh*Dmrt2* electroporated cells. Eventually, post-mitotic cells will not be generated to populate the CP, which is what was observed. **4)** Most GFP(+) cells extend projections to the thalamus, similar to mock brains. However, a subset of projections is misguided laterally, above the external capsule, instead of following the corticothalamic track. In the sh*Dmrt2* brains, this tract and the cc are de-fasciculated (**Figure 4a’’’-d’’’; Table 2; Suppl. Table 1, 2**). In summary, these results demonstrate that *Dmrt2* downregulation affects the neurogenesis and proper development of CgCx deeper neurons.

**Figure 4.**
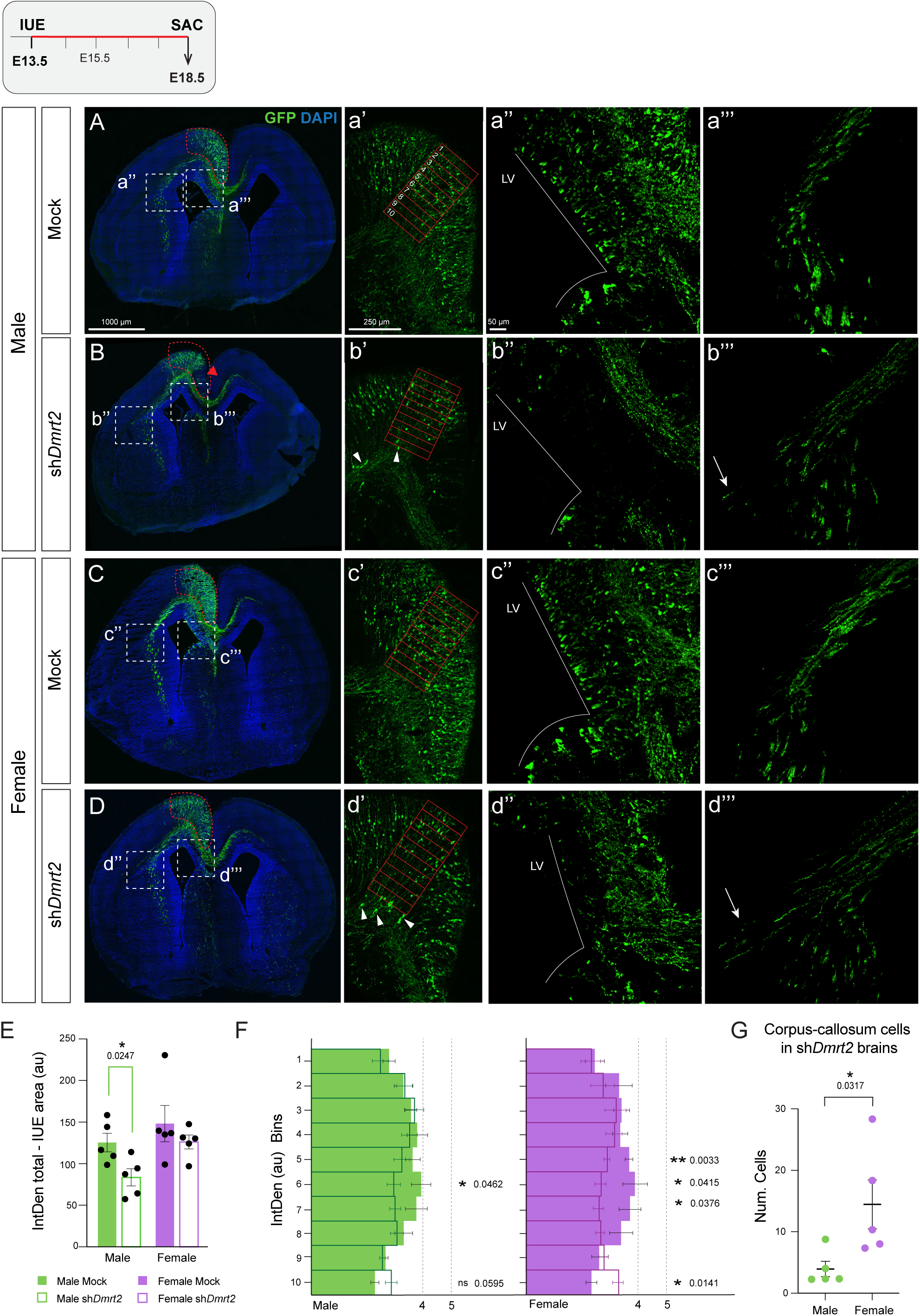
*Dmrt2* downregulation alters the development of cingulate cortical neurons. **A-D)** Representative images of E18.5-electroporated brains at E13.5. **A) Male mock, B) Male sh*Dmrt2*, C) Female mock, and D) Female sh*Dmrt2*.** The electroporated (red dashed circled) area is quantified in E). **a’-d’)** Higher magnification of the cingulate cortex in the four conditions. The red box area was divided into 10 bins to cover the cortical plate of the CgCx and fluorescence was quantified in F). White arrowheads indicate ectopic GFP(+) cells in the cc of sh*Dmrt2*-treated animals, quantified in G. **a’’-d’’**) Higher magnification for the ventricular zone (VZ). In *shDmrt2* animals, GFP(+) cells are not visible in this area. The white line delimits the lateral ventricle (LV). **a’’’-d’’’**). Higher magnification of the GFP(+) projections towards the thalamus across the cc, above the internal capsule. White arrows indicate lateral misguidance. **E) Decrease in the total GFP fluorescence intensity in male embryos**. The IUE area was quantified by measuring the GFP fluorescence, which was determined by integrated density (IntDen). Two-tailed t-test (n = 5). **F) Cellular distribution of GFP(+) cells in the medial CgCx.** The histogram represents each bin’s GFP fluorescence intensity (IntDen), marked in a’-d’. Solid bars depict mock embryos, while empty bars represent *shDmrt2*-treated animals. Two-tailed t-test (n ≥ 5). **G) Count of the GFP(+) staked cells in sh*Dmrt2* brains.** Two-tailed t-test (n = 5). In scatter plots, each dot represents individual embryos. In all graphs, the results correspond to the mean±SEM of the average of three sections per embryo, and the significance is as follows: * p-value < 0.05 and ** p-value < 0.01.

Males had a more pronounced depletion of GFP-positive neurons, while in females, the remaining sh*Dmrt2* electroporated cells presented exacerbated phenotypes compared to males and mock cells. To investigate this apparent contradiction, we conducted a transcriptional profile of electroporated cells in both sexes in mock and downregulated conditions.

### Loss of general neuronal identity of CgCx neurons upon *Dmrt2* depletion

Based on our anatomical analysis, *Dmrt2* downregulation affects the correct specification of cingulate neurons by controlling their migration and targeting. We asked whether *Dmrt2* could have a role in acquiring neuronal terminal identity. For this experiment, we performed an unbiased RNA-Seq from fluorescence-activated cell-sorted (FACS) electroporated cells. Like the previous experiment, embryos were electroporated at E13.5 into the CgCx primordium. At E18.5, FACSorted electroporated fluorescence cells were selected, and total mRNA was isolated and sent for sequencing (see **Experimental Procedures** for details).

The principal component analysis (PCA) showed that sh*Dmrt2*-treated embryos are clustered together, separate from mock embryos. Within sh*Dmrt2* samples, male and female embryos are mixed and do not segregate by sex, while mock samples are further clustered by sex (**Figure 5A**).

**Figure 5.**
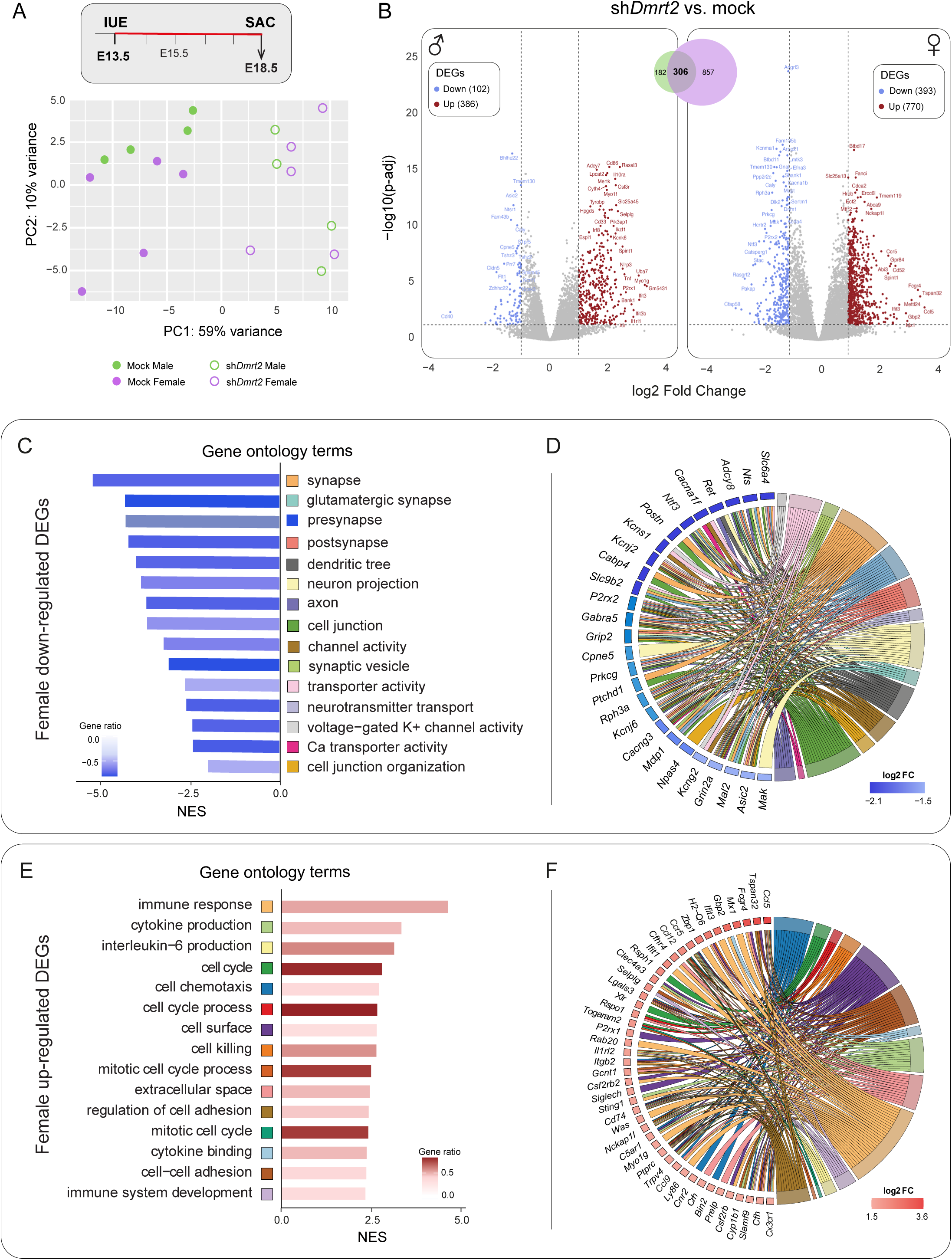
*Dmrt2* controls neuronal differentiation in post-mitotic neurons. **A)** Two-dimensional PCA plot for the four experimental conditions shows the clustering of mock separated from sh*Dmrt2* samples. Among the sh*Dmrt2* group, male and female samples show less clear clustering than in mock conditions. **B)** Volcano plots represent DEGs in sh*Dmrt2* vs. mock comparisons (**Suppl. Tables 3 and 4**). Blue indicates downregulated genes, and red indicates upregulated genes (numbers in brackets). More DEGs are retrieved in female comparisons (1163) than male ones (488). The trend in down- and up-regulated DEGs is similar in the two sexes. The Venn Diagram indicates the number of DEGs shared (306) by male and female comparisons. **C-F) Female sh*Dmrt2* vs. mock DEGs analysis. C)** GSEA on down-regulated DEGs shows neuron-related overrepresented categories. 15 categories with NES < 2 (out of 136; **Suppl. Table 3**) are shown in the bar plot. **D)** Among the 15 down-regulated categories, genes with changes between log2FC of −2.13 and −1.51 are represented in the GO-Chord plot. **E)** GSEA on up-regulated DEGs are related to immune system and cell cycle. 15 categories with a NES > 2 (out of 159, **Suppl. Table 3**) are shown in the bar plot. **F)** Among the 15 up-regulated categories, genes with changes between log2FC of 1.5 and 3.5 are represented in the GO-Chord plot.

To proceed with the analysis, we selected Differentially Expressed Genes (DEGs) with an adjusted p-value (p-adj) < 0.05 and a log2 Fold Change (FC) ≥ |1| (only genes that are at least two-fold up- or down-regulated) between four comparisons: mock male vs. mock female; sh*Dmrt2* male vs. sh*Dmrt2* female; sh*Dmrt2* male vs. mock male and, lastly, sh*Dmrt2* female vs. mock female (**Figure 5B**; **Suppl. Figure 5B; Suppl. Tables 3-5**). Overall, in the male-to-female comparisons, both in mock and sh*Dmrt2*, only a very reduced number of DEGs were retrieved (3 genes in mock: *Pf4* is downregulated and *Scml4* and *Serpina3n* are upregulated; 6 genes in sh*Dmrt2* comparisons: *Stk11*, *Npm3*, *Cd151* and *Mpst* are downregulated, while *Lhx6* and *Stac* are upregulated (**Suppl. Figure 5B**). However, when we compared sh*Dmrt2* vs. mock-treated cells in each sex, the differences in the number of DEGs were more pronounced. We retrieved 2.4 times more DEGs in female than male comparisons (**Figure 5B**).

Our initial analysis focused on *Dmrt2* expression levels. Surprisingly, in male cells, we observed that the sh*Dmrt2* treatment did not result in a statistically significant reduction of *Dmrt2* expression compared to the mock treatment (15.5% reduction; p-adj = 0.4524) (**Suppl. Figure 5C**; **Suppl. Table 4**). On the contrary, females show a 56.25% reduction (p-adj = 0.0216) (**Suppl. Table 3**). We then conducted a comparison between male and female datasets, which yielded 488 and 1163 DEGs, respectively, and found that approximately two-thirds of the DEGs identified in males were also present in the female dataset, indicating a substantial overlap in gene expression changes between sexes (**Venn diagram in Figure 5B; Suppl. Table 5**).

To comprehensively characterize the phenotype, we conducted a gene ontology (GO) study by Gene Set Enrichment Analysis (GSEA) on sh*Dmrt2* vs. mock DEGs. **Figure 5C-5F** shows only DEGs from female comparisons.

On the female comparison GSEA, we selected 15 categories with Normalized Enrichment Scores (NES) > |2| (**Figure 5C, 5D**; **Suppl. Table 3**). The analysis of the most significant downregulated categories suggests that migration and neuronal functions are altered. Additionally, cell junction organization, calcium (Ca) and potassium (K) channel activity genes [Ca-channel subunits (*Cacna1f*, *Cacng3*) and Ca binding proteins (*Cabp4*, *Cpne5, Rph3a*), K-channel subunits (*Kcns1*, *Kcnj2, Kcnj6, Kcng2*)], neurotransmitter pathway genes [serotonin transporter *Slc6a4*/*Sert* and glutamate receptors and interacting proteins *Grin2a, Grip2*], synaptic vesicle, pre and post-synapse activity [the transmembrane protein, such as *Ptchd1*, *Npas4* or *Prkcg/PKC-gamma*] show the highest reduction in expression in the female comparison.

The most significant GO terms among the up-regulated DEGs include immune response, cytokine production, cell cycle, cell chemotaxis, cell killing, extracellular space, and regulation of cell adhesion (**Figure 5E, 5F**). *Tumor necrosis factor (Tnf)* and several genes encoding TNF receptors (*Tnfrsf1b, Tnfrsf13b*…, among others) or TNF-related proteins (*Tnfaip8*) are also upregulated. The TNF pathway has been involved in neurite outgrowth inhibition^30^. Genes involved in the secretion of extracellular matrix components are also upregulated (several matrix metallopeptidases or *Mmp* genes or collagens like *Col24a1,* for example)^31^. *Dmrt2* regulates (directly or indirectly) many genes that encode cell adhesion molecules (CAMs), such as cadherins, nectins, and netrins, that are required in the control of many neurodevelopmental aspects, including neural migration, axonal pathfinding, and wiring, along with synapse formation^32^. These processes regulate neurogenesis, cell migration, and axonal outgrowth and guidance^30–35^. Many genes encoding chemokines and cytokines are among the most up-regulated genes [*Ccl5*, *Ccl12*, *Ccl9,* among others] and their receptors [*Ccr5* and *Cx3cr1*]. Canonical markers of microglia like *Cx3cr1* and *Iba1* (also known as *Aif1*) show up in our list of DEGs. We discarded the possibility of microglia contamination in our FACSorted cells because only electroporated ventricular progenitors carry the *pDsRed* plasmid.

Most *Dmrt2* neurons in the CgCx can be identified as TBR1-CThPN. However, *Dmrt2* does not regulate any of the transcription factors that have been previously characterized for cortical projection neurons subtype regulators, except for *Satb2* expression, which is reduced at approximately 50% of its level in mock conditions (log2FC = −0.97; p-adj = 0.00013), and *Ctip1* (also known as *Bcl11a*), which is moderately reduced by 28% (log2FC = −0.48; p-adj = 0.01). Notably, the expression of *Tbr2* (also known as *Eomes*) showed a 2-fold increase in *Dmrt2*-depleted neurons (log2FC = 1.07; p-adj = 1.45e-10). This substantial upregulation, coupled with a moderate rise in *Pax6* expression (log2FC = 0.73, p-adj = 0.00026) and increased levels of cell cycle-related genes (**Figure 5E**), indicates that neurons with reduced *Dmrt2* expression may retain characteristics of an immature state.

The GSEA analysis of all male genes also shows terms related to neuronal function in the downregulated set and the immune system in the upregulated one. Although similar biological processes are affected in both sexes, a few categories were specifically found in the male comparisons (**Suppl. Figure 5D, 5E**) related to olfactory receptor activity or sensory perception of smell [vomeronasal 2 receptor genes (*Vmn2r3* and *Vmn2r82*) or several olfactory receptors (*Or2w3b* and *Or8b1* among others)], foregut morphogenesis or metabolic processes such as tetrapyrrole biosynthetic process. Although all these genes show significant p-values in female comparison, they do not meet the p-adj criteria. Most likely, the same processes are affected in both sexes but with differential magnitude (**Suppl. Table 4**).

In conclusion, the gene expression profile of cells with reduced *Dmrt2* levels reveals a complex pattern of transcriptional changes. Specifically, we observed a decrease in the expression of neuron-specific genes and an increase in non-neuronal gene expression. Most genes regulated by *Dmrt2* are functional effectors and not master regulators of cortical projection neuron identities. This transcriptional shift suggests that *Dmrt2* plays a crucial role in terminally defining the overall neuronal identity of CgCx neurons.

## DISCUSSION

We have uncovered a novel function for the transcription factor *Dmrt2* in mouse cortical development. Specifically, we show an unprecedented function in the coordination of cingulate cortex neuronal differentiation. Our research deepens our understanding of the role of *Dmrt2*, going beyond its established peripheral functions. It also addresses a broader question regarding the interaction between genetic factors that regulate sexual differentiation and those that shape general brain development in mammals. Furthermore, we explored the conservation of *Dmrt* transcription factors in the sexual differentiation of the mammalian nervous system and identified sex-specific susceptibilities between male and female embryos upon *Dmrt2* downregulation.

### *Dmrt2.1* presents sexual differences in levels of expression during cingulate cortical development

Our results show that *Dmrt2* is highly enriched in post-mitotic CgCx neurons and expressed in deep-layer neurons across the entire rostrocaudal extension of the CgCx. Expression in this region is maintained from the early embryo until adulthood. This result, together with the previous report from Galazo *et al.*^15^, situates *Dmrt2* in CThPNs. Our analysis further shows that *Dmrt2* is not restricted to CThPNs, marked by TBR1, but a few scattered *Dmrt2* neurons colocalize with SATB2 in potentially callosal neurons. Additional tracing studies should be performed to confirm specific projections that might extend from *Dmrt2*-neurons to locations other than the thalamus.

At least three predicted *Dmrt2* variants will be transcribed from the *Dmrt2* locus (NCBI Gene ID: 226049) due to exon 4 retention, skipping, or transcription initiation. However, the only variant that showed statistically significant sexual differences was *Dmrt2.1,* while *Dmrt2.2* and *Dmrt2.3* did not show any differences at any time point. The lack of spatial resolution hampered our ability to discern whether the origin of the sexual difference comes from different levels in all *Dmrt2*-expressing cells or expression in distinct cell types. Single-cell RNA-Seq experiments comparing male and female embryos could shed light on this question, but unfortunately, such information is not available for the cingulate cortex.

Although no function has been assigned yet to *Dmrt2* alternative variants, future studies should be performed to uncover additional layers of regulation. DMRT2.1 and DMRT2.2 protein products differ in their C-terminal ends and could be controlling different target genes by binding distinct cofactors. These two variants also differ in their 3’ untranslated regions where post-transcriptional regulation involving microRNAs is crucial for controlling gene expression. Indeed, *in vitro* studies in neuroblastoma cell lines have shown that human *DMRT2* expression is regulated by *miR-16-5p*^24^. Additionally, post-transcriptional interactions between variants might exist. The potential product of *Dmrt2.3* bears no DNA binding domain and could act as a dominant negative. The human *DMRT2* locus is more complex, and many more variants are predicted to be transcribed (NCBI Gene ID: 10655 and^36^). However, a striking level of conservation in the regulatory logic between the human and murine loci makes our findings relevant for human studies. In humans, eleven predicted *DMRT2* mRNA isoforms involve exon 4 retention or skipping, a process that in mice produces *Dmrt2.1* and *Dmrt2.2*, and alternative transcription initiation.

Interestingly, the sexual difference in *Dmrt2.1* levels was observed at E13.5 before the peak of testosterone secretion from the embryonic gonad had occurred yet^37^. Therefore, *Dmrt2.1* transcriptional differential levels might have an intrinsic chromosomal origin. DMRT TFs have been shown to control somatic morphological dimorphisms across phylogeny^4,5,7,38^, which is usually ensured by their binary dimorphic expression. In the case of CgCx, we did not observe a dimorphic expression of *Dmrt2*; rather, we found quantitative differences between the sexes. Although subtle, these early differences in expression can lead to significant functional consequences, resulting in distinct developmental outcomes between males and females at later stages. It is conceivable that as development progresses, we would have observed even more profound sexual functional differences in postnatal animals. Still, for those types of experiments, it is necessary to use more robust genetic models like conditional *Dmrt2* knock-outs where spatial and temporal manipulations could be performed, and rescue experiments could be done.

### *Dmrt2* maintains the proliferative capacity of cortical progenitors, either through direct regulation or indirect mechanisms

*Dmrt2* downregulation triggers the decrease in progenitor cells within the ventricular zone of the cingulate primordium. Conversely, *Dmrt2.1* overexpression produces the opposite outcome, increasing the ventricular zone proportion of cells at the expense of generating post-mitotic neurons. This result aligns with a previous report on human neuroblastoma cell lines where DMRT2 positively regulates the proliferation rate and progression of tumor growth^24^. ISH techniques employed in this study failed to detect *Dmrt2* mRNA expression in cortical progenitor cells. Furthermore, the absence of commercially available antibodies precludes direct examination of DMRT2 protein distribution, which will eventually determine the TF function. We can envision two possibilities to explain the effect on progenitors: 1) a non-cell autonomous role for *Dmrt2* in the control of proliferation of progenitor cell cycle. Many genes we found misexpressed in *Dmrt2* downregulation cells encode for secreted molecules that might impact progenitors, or 2) *Dmrt2* expression levels are below the ISH detection threshold. The fact that we detect expression with qPCR at E13.5 before we can observe any expression in the cingulate primordium through ISH, including in the CP where *Dmrt2* is clearly detected from E14.5, opens the possibility that *Dmrt2* is not detected in the VZ at any time point (not at E13.5 nor later) with this technique. Transcription factors often function at very low concentrations^39^. However, subtle changes in transcription factor levels can have significant functional consequences^40^.

Other *Dmrts* genes, the DMA subfamily, have been previously described as having a role in the proliferation of cortical progenitor cells in mice. *Dmrt3* and *Dmrt5* are required to maintain the neural precursor cell proliferation in the medial pallium, which will generate the hippocampus^11^. This role is extensive to other species and has been extensively studied in the NS of *Drosophila,* where *doublesex* regulates neuroblast proliferation in a sex-specific manner^41–43^. Therefore, the interaction of *Dmrts* with the cell cycle becomes a recurring theme, and specific mechanistic details in neurons remain to be explored.

### *Dmrt2* controls general aspects of neuronal differentiation in post-mitotic neurons, including migration, axonal targeting, and gene expression

*Dmrt2* has been previously described as an activator in the later stages of myoblast and chondrocyte differentiation. It is involved in the balance between self-renewal and differentiation^16,17^. Our research reveals that *Dmrt2* acts as a transcriptional regulator of the general terminal identity of deeper-layer neurons of the cingulate cortex.

We observed no changes in the expression of CThPN master regulators such as *Tbr1* or *Fog2*, suggesting that *Dmrt2* may operate downstream of this regulatory network. Galazo *et al*., proposed *Dmrt2* to be a late developmental gene that might be downstream of FOG2, although this was not tested specifically in the study^15^. Despite the limitations of our RNA-Seq in bulk approach, which precluded analysis of cell-specific processes, the observed reduction (although modest) in *Ctip1* and *Satb2* expression could indicate that one of the most affected neuron types are the deeper callosal neurons of layer V. It has been reported that in the absence of CTIP1 function, more subcortical projection neurons (SCPNs) are generated at the expense of both CThPNs and deeper CPNs^44^. SABT2 induces CPN identity and SCPN identity^45^ with a limited role in regulating CThPN axons^26^. However, these TFs have been proposed to act in a context-dependent manner and have been mostly studied outside the CgCx, where layer and neuronal specification are not so well studied. It remains to be determined, with cellular resolution, the epistatic relationships between *Dmrt2* and them.

Numerous target genes involved in terminal neuronal functions have been identified to be regulated, directly or indirectly, by *Dmrt2*. The most overrepresented downregulated GSEA ontologies are fundamental for neuronal circuit assemblies and communication. Conversely, the upregulated-transcriptional landscape of *Dmrt2*-depleted cells reveals characteristics of an undifferentiated state (such as the upregulation of *Tbr2* and *Pax6*, or chemokine receptors, reported to be expressed in cortical progenitors^46^) and the inability to acquire the terminal identity of postmitotic projection neurons. This pattern of transcriptional changes suggests that *Dmrt2* may play a complex regulatory role in balancing neuronal specification processes during late-stage differentiation.

One of the most prominent phenotypes upon *Dmrt2* depletion is the impaired migration of deeper layer neurons and misguided projections and de-fasciculation. The RNA-Seq *Dmrt2*-regulated targets corroborate the dysregulation of pathways governing these developmental processes: *Ctip1*, for which we found a modest reduction, is known to regulate these processes, and aberrant commissural projections are found in the CgCx of *Ctip1^Flox^;Emx^Cre^* conditional knock-out mice^47^; or CAMs required in the control of many neurodevelopmental aspects, including neural migration, axonal pathfinding, and wiring, along with synapse formation are also affected^32,33,48^. Moreover, the prominent upregulation of chemokine-encoding genes and their receptors might directly affect neuronal migration defects. Neuronal expression of chemokines has been described before, and their downregulation in neurons has been proposed to influence many aspects of neuronal development, including differentiation, survival, and synaptic transmission^46,49–51^.

### *Dmrt2* may interact with intrinsic sex factors

By downregulating *Dmrt2* expression when deeper cortical neurons are born, we have uncovered potential sex differences in *Dmrt2*-dependent function.

Proliferation rates are not distinct between the sexes in mock embryos. However, when *Dmrt2* is depleted, latent sexual differences emerge, bearing males stronger phenotypes in progenitors. *Dmrt2* downregulation unmasks sex-specific differences, prompting male progenitors to exit the cell cycle more rapidly than female ones and progress toward differentiation. This topic requires more research, but interestingly, glioblastoma, the most common brain tumor, presents sexual differences in the incidence and outcome. Such sex differences exist across all stages of life, indicating some independence from hormone action^52^. Although the precise mechanisms are unknown, a clinical correlation was found between cell cycle and integrin signaling.

The transcriptional profile analysis in late embryos reveals seemingly contradictory results, with female comparisons exhibiting higher numbers of DEGs than male comparisons, although male progenitors are more substantially affected. We interpret these results as a consequence of a more substantial depletion of the male progenitor pool that yields fewer neurons, where the shRNA would result in higher concentrations per cell than female sh*Dmrt2*-treated cells. Cells with higher shRNA levels could exhibit a compensatory “rebound effect”, only observable in male cells, restoring *Dmrt2* expression to mock-equivalent levels and producing milder phenotypes in late male embryos. This compensation aligns with prior studies that demonstrate that high siRNA concentrations accelerate and amplify rebound responses in both *in vitro* and *in vivo* models to a greater extent^53,54^.

### *DMRT2* and human intellectual disability

The CgCx is anatomically and functionally connected to a broad set of regions engaged in social information processing. Therefore, genetic mutations affecting its development could be associated with neurodevelopmental disorders such as cognitive disability, autism spectrum disorders, epilepsy, schizophrenia, and attention deficit hyperactivity disorder^55^. In this study, we report that *Dmrt2* regulates the proliferation and specification of deeper-layer CgCx projection neurons and that its effects are aggravated in male mice. The cingulate cortex has been postulated as one of the brain areas that shows sex differences in the elicited response to emotional stimuli^56^ or anatomical features^57^. In rodents, sex differences have been found at the synaptic and connectivity levels^58,59^. Notably, it has been reported that males have a higher proportion of layer V neurons in the CgCx projecting to the prefrontal cortex than females^58^. Additional sex differences have been found at the level of synaptic plasticity in CgCx^59^. If similar mechanisms to those described in this paper are conserved, *DMRT2* could be involved in the development of the projection neurons of the human cingulate cortex. This could potentially explain males’ increased vulnerability to neurodevelopmental disorders, providing a molecular basis for observed sex differences in these conditions. According to the Human Protein Atlas database, DMRT2 is present in the brain (https://www.proteinatlas.org/), but no other reports document its expression in the human brain.

*DMRT2* is close to *DMRT1* and *DMRT3* in the short arm of chromosome 9. Deletions or duplications in that chromosomal region have been associated with intellectual disability and mental disorders^19,21,22,60^. *DMRT1* has been extensively studied in the human and mouse gonad^5^, and no apparent neuronal expression has been reported. In mice, the DMA-DMRT *Dmrt3* is expressed in cortical progenitors^9^, hindbrain^61^, and spinal cord^62^. Potentially, its deletion could be related to intellectual disability in humans as well. However, this has not been proven yet, and no behavioral studies have been performed in mouse *Dmrt3* mutants or human patients with *DMRT3* mutations. Whether *DMRT2* mutations contribute to human intellectual disabilities through an aberrant development of the cingulate cortex will be of significant interest to investigate further.

## EXPERIMENTAL METHODS

### Animals and tissue preparation

Animals used in this study include male and female C57BL/6JRccHsd (ENVIGO) and C57BL/6J (The Jackson Laboratory) wildtype mice. Animals were housed and bred at the CBM animal facility and treated according to European Communities Council Directive 86/609/EEC regulating animal research. All procedures were approved by the Bioethics Subcommittee of Consejo Superior de Investigaciones Científicas (CSIC, Madrid, Spain) and the Comunidad de Madrid (CAM) under the following protocol approval number (PROEX 286/19; RD 53/2013). The presence of a vaginal plug estimated embryonic day (E) 0.5 of pregnancy after overnight male mating. Embryos were collected from independent litters.

For embryonic sex genotyping, tail tissue was collected and digested in 50 μl of MgCl_2_-free PCR reaction buffer (10013-4104, Biotools) supplemented with 400 μg/ml proteinase K (3115879001, Roche) at 55°C with shaking for 1-3 hours (h), followed by heat-inactivation of the proteinase K at 95°C for 5 min. One microliter of a 1/10 dilution of each sample was added directly to a 25 μl PCR reaction. The protocol of genomic DNA amplification was adapted from McFarlane *et al.*^63^. The primers used were: *Sx*-Forward 5’-*GATGATTTGAGTGGAAATGTGAGGTA*-3’ and *Sx*-Reverse 5’-*CTTATGTTTATAGGCATGCACCATGTA*-3’ and the thermal protocol was: initial cycle of 5 minutes (min) at 94°C followed by 35 cycles of 45 seconds (s) at 94°C denaturalization, 30 s at 55°C annealing and 1 min at 72°C extension with a final extension step at 72°C for 10 min.

For staining, brains from embryos (E13.5, E14.5, E15.5, and E18.5) were fixed in 4% paraformaldehyde (PFA) (1.04005.1000, Merck) diluted in phosphate buffer (PBS 1X) for 2 h. Postnatal (P) 7 brains were fixed in 4% PFA for 4 h at room temperature (RT). Adult brains (P56) were fixed in 4% PFA overnight (o/n) at 4°C from animals previously perfused intracardially with sterile PBS 1X followed by 4% PFA. After fixation, brains were washed in PBS 1X several times, and cryoprotected in 15% sucrose (1.07651.1000, Merck)/PBS 1X. Lastly, they were embedded in 7.5% gelatin (G2625, Sigma)/15% sucrose (1.07651.1000, Merck)/PBS 1X and frozen at −80°C. Coronal cryostat sections were cut at 16 μm thickness.

For total RNA isolation, tissue was freshly collected from embryo and postnatal mouse brains and homogenized with 3 mm stainless steel Lysing beads (Alpha Nanotech, VWR) in diethyl pyrocarbonate (DEPC)-treated PBS (D5758, Sigma-Aldrich) for 1 min at 30 Hz with a TissueLyser (MM300, Retsch). Fresh or homogenized tissue was stored at −80° until use.

### *In situ* hybridization (ISH)

ISH was carried out as previously described by Di Meglio *et al.*^64^. Briefly, brain sections were incubated with 4% PFA for 10 min at 4°C. Then, prehybridization was performed at RT with hybridization buffer [50% deionized formamide (S4117, Millipore), SALTS 1X (Tris 10 mM, NaCl 200 mM, NaH_2_PO_4_ 5 mM, Na_2_HPO_4_ 5 mM, EDTA 5 mM), Denhardt’s 1X (D2532, Sigma), 10% dextran sulphate (4911, Sigma), tRNA 1 mg/ml (R6625, Sigma, Merck)] for 1 h in a humified chamber with 5X SSC [sodium citrate 750 mM, NaCl 75 mM] and 50% deionized formamide (S4117, Millipore). Tissue sections were incubated with the anti-sense digoxigenin-labeled (11277073910, Roche) probe (1:1000 in hybridization buffer) o/n at 72°C. Following hybridization, the slides were washed in SSC 0.2X for 90 minutes at 72°C, blocked in 2% blocking solution (11096176001, Roche) in MABT at pH 7.5 [maleic acid 100 mM, NaOH 200 mM, NaCl 200 mM, 0.1% Tween-20 (P1379, Sigma-Aldrich)] for 1 h at RT and then incubated o/n at RT with anti-digoxigenin-alkaline phosphatase antibody (11093274910, Roche) at 1:5000 diluted in MABT. After several washes, the alkaline phosphatase activity was developed using 3.5 μl/ml BCIP (11383213001, Roche) and 1 μl/ml NBT (11383221001, Roche) diluted in NTMT solution at pH 9.5 [Tris 100 mM, NaCl 100 mM, MgCl_2_ 50 mM, 0.1% Tween-20 (P1379, Sigma-Aldrich)] for 20 h at RT. *Dmrt2* probe covers the whole gene except the first ATG and was kindly provided by Takahiko Sato^16^ (see **Figure 1E**).

### Fluorescent *in situ* hybridization (FISH)

The initial steps of hybridization were performed as described above. For detection, horseradish peroxidase-conjugated anti-digoxigenin antibody at 1:5000 (11207733910, Roche), TSA™ Plus Biotin System (NEL749A001KT, Perkin Elmer®), and 1:250 Streptavidin-Alexa 555 (S32355, Thermo Fisher) were used according to the manufacturer’s instructions.

### Immunofluorescent staining (IF)

IF was performed following standard protocols. Brain sections were washed with PBS 1X with 0.1% Triton™ X-100 (T9284, Sigma-Aldrich) (PBST) and blocked 1 h in blocking solution [PBST and 1% of bovine serum albumin (BSA) (A2153-100G, Sigma)]. The following primary antibodies diluted in a 1/10 blocking solution were incubated o/n at 4°C: chicken anti-GFP 1:1000 (ab13970, Abcam), rabbit anti-TBR1 1:300 (ab31940, Abcam), mouse anti-SABT2 1:50 (ab51502, Abcam), rat anti-CTIP2 1:500 (ab18465, Abcam), rabbit anti-Ki67 1:1000 (ab15580, Abcam) and sheep anti-TBR2 1:200 (AF6166, R&D System). In the case of TBR2, heat-induced antigen retrieval was performed before primary antibody incubation (citrate buffer 10 mM, pH6, for 5 min at 110°C in a boiling chamber). Corresponding secondary antibodies were incubated at 1:1000 for 2 h at RT: goat anti-chicken-Alexa 488 (A11039, Thermo Fisher), donkey anti-mouse-Alexa 555 (A31570, Thermo Fisher), donkey anti-rabbit-Alexa 555 (A31572, Thermo Fisher), donkey anti-rabbit-Alexa 647 (A31573, Thermo Fisher), goat anti-rat-Alexa 555 (A21434, Thermo Fisher) and donkey anti-sheep-Cy3 (713-165-003, Jackson ImmunoResearch). Hoechst 33342 (H1399, Invitrogen™) was used for cell nuclei staining.

For double ISH or FISH and IF staining, ISH was carried out prior to IF as previously described^64^.

### Imaging

ISH chromogenic and IF images were obtained with a DMCTR5000 microscope equipped with DFC500 color and DFC350 FX monochrome cameras (Leica), respectively.

Tilescan mosaic images were acquired by SpinSR10 disk confocal system with an IX83 inverted microscope (Olympus) and reconstructed with CellSens Dimension 4.2.1 software (Olympus). All images were acquired using a 1024 × 1024 scan format with a 40x or 60x objective.

Integrated density (IntDen) was calculated using Fiji. For total IntDen, the IUE area was limited to the cingulate cortex and outlined manually. For cellular distribution, IntDen was measured in a defined area spanning from the cortical layer I to the corpus callosum (not included), subdivided into 10 equal bins.

### Quantitative real-time PCR (qPCR)

Total RNA was isolated using NZY Total RNA Isolation kit (MB13402, Nzytech) following the manufacturer’s instructions. cDNA was obtained from 1 μg of total RNA with First-Strand cDNA Synthesis kit (27-9261-01, GE) in a 15 μl reaction volume. qPCR was performed using GoTaq® qPCR Master Mix (A6002, Promega) following the protocol of the manufacturer in a CFX-384 Touch Real-Time PCR Detection System (Bio-Rad). The following gene-specific primer pairs were used: *Dmrt2.1*-Forward and *Dmrt2.2/2.3*-Forward 5’-*AGGGGCTTTCTGGGAAACAG*-3’, *Dmrt2.1*-Reverse 5’- *CGTCATTGGGCGATAACCTTC*-3’, *Dmrt2.2/2.3*-Reverse 5’- *TAGAATAATCCAAGGAGAACTTC*-3’, *Dmrt2-Ex1*-Forward 5’- *CCTGAGCCTGGTTCTTG*-3’, *Dmrt2-Ex1*-Reverse 5’- *CAGTGGAGTCCCACAGCTATC*-3’, *GFP*-Forward 5’-*CAACCACTACCTGAGCACCC*- 3’, *GFP*-Reverse 5’-*GTCCATGCCGAGAGTGATCC*-3’, *Gus*-Forward 5’- *AGCCGCTACGGGAGTCG*-3’ and *Gus*-Reverse 5’-GCTGCTTCTTGGGTGATGTCA-3’. *Dmrt2* and *GFP* expression levels were quantified in triplicate and normalized to *Gus* expression levels. The data were analyzed using the comparative Ct method.

### Retro-transcription PCR (RT-PCR)

For the end-point RT-PCR, 400 ng of the qPCR-extracted total RNA were reverse-transcribed using SuperScript™ III One-Step RT-PCR System with Platinum™ *Taq* High Fidelity DNA Polymerase (12574035, Invitrogen™) according to the manufacturer’s recommendations in a 20 μl reaction volume.

The following cycling conditions were established using a T100 Thermal Cycler (Bio-Rad): initial cycle of 30 min at 55°C and 2 min at 94°C followed by 40 cycles of 15 s at 94°C denaturalization, 30 s at 58°C annealing and 30 s at 68°C extension with a final extension step at 68°C for 5 min. *Hprt1* was amplified as loading control. Primers used in this study were as follow: *Dmrt2-Ex4*-Forward 5’-CGGAAAGCAGTGTACCAGAG-3’, *Dmrt2-Ex4*-Reverse 5’-CTTGTCAGCAAAGGCTCTTC-3’, *Hprt1*-Forward 5’-TCCTCCTCAGACCGCTTTT-3’ and *Hprt1*-Reverse 5’-CCTGGTTCATCATCGCTAATC-3’.

### *In utero* electroporation (IUE) and plasmids

IUE was performed as previously described in Briz *et al*.^65^. Briefly, the mixture composed of: a plasmid expressing GFP or DsRed fluorescent protein (1 μg/μl), 0.1% Fast Green FCF (F7252, Sigma) and, in the case of silencing or overexpression experiments, sh*Dmrt2* or *pCAG*-*Dmrt2.1* plasmid (1 μg/μl) respectively, was injected into the left lateral ventricle of E13.5 embryos using pulled glass micropipettes. Five voltage pulses (40 mv, 50 ms) were applied using external paddles oriented to target the cingulate cortex. By E15.5 and E18.5, GFP- or Red-brain embryos were selected for further analyses.

Plasmids used were *pCAG-GFP* (#11150, Addgene); *pCAGGs-DsRedExpress* (gift from Marta Nieto); *Dmrt2* short hairpins, sh*Dmrt2*_1 and sh*Dmrt2*_2, in pLKO.1 vector were acquired in Sigma-Aldrich [hairpin sequences are sh*Dmrt2*_1: *AGCAGACGCTCAGCGACAAAT*(TRCN0000418486), sh*Dmrt2*_2: *GCCACAACTTACAGACAATAT* (TRCN0000086379) and, sh*Dmrt2*_3: *GCTTACAAGGGAAGCTGGATT* (TRCN0000086378)] (see **Suppl. Figure 2A** for approximate location of each sh in the *Dmrt2* locus).

*Dmrt2.1* cDNA was cloned into the pCAG backbone obtained from the double-digested *pCAG-GFP* plasmid (#11150, Addgene) by EcoRI (10703737001, Roche) and NotI (11037668001, Roche). cDNA was isolated from the *Dmrt2*-probe plasmid, kindly provided by Takahiko Sato^16^. The primers used were: Forward 5’- *TATCGAATTCTGGCCACCATGACGGAAGGGCAGGCAG*-3’, including EcoRI-recognition and Kozak sequences, and Reverse 5’- *GATTGCGGCCGCTTTACTGAGAGACGTGAGTGAC*-3’, including NotI-recognition sequence. The ligation reaction was carried out using the DNA ligase T4 (M0202L, New England BioLabs) in T4 ligase buffer (B0202S, New England Biolabs) according to the manufacturer’s specifications, o/n at 16°C.

To investigate proliferation of neural progenitor cells in electroporated embryos, intraperitoneal injection with 5 mg/kg 5-ethynyl 2’-deoxyuridine (EdU) (C10339, Invitrogen™) were given into pregnant mice 2 h before harvesting. Then, the embryonic brains were collected following the staining protocol above described. For EdU labelling detection, Click-iT™ EdU Cell Proliferation kit for imaging, Alexa Fluor™ 594 dye (C10339, Invitrogen™) was used following the manufacturer’s instructions.

### RNA-Sequencing and FACSorting

For the RNA-Seq, embryos were electroporated with *pCGA-DSRedExpress* and sh*Dmrt2* as previously described. At E18.5, individual embryos with fluorescence signal were selected, and brains were collected on ice-cold D-PBS (14287-072, Gibco™). After meninges removal, cingulate cortices were isolated, finely chopped and digested for 15 min in D-PBS containing Liberase 0.15 mg/ml (05 401 119 001, Roche), 300 U of DNase I (18047019, Invitrogen™). After addition of D-PBS-/0.05% Fetal Bovine Serum (FBS) (A5256701, Gibco), the tissue was mechanically dissociated with 230 mm-length glass Pasteur pipettes (702, Deltalab), whose tips have been previously fire-polished reduced five times. Cell debris was eliminated using Debris Removal Solution (130-109-398, Miltenyi Biotec) following the manufacturer’s specifications and the pellet was recovered. Cells were resuspended in sorting medium [0,5% FBS, trehalose 0.05 mg/ml (T9531, Sigma) and in D-PBS] and treated with 100 U of DNase I (18047019, Invitrogen™). Samples were not pooled and each sample corresponded to one single DsRed(+)mouse brain. Lastly, cells were sorted using FACSAria™ Fusion Cell Sorter (BD Biosciences). Cell size, complexity and fluorescence intensity were acquired with BD FACSDiva™ 8.0 Software. Analysis was performed using FlowJo™ v10.9 Software (BD Life Sciences).

RNA was extracted and purified from the sorted cell suspension with RNeasy® Plus Micro Kit (74034, Qiagen) following the manufacturer’s protocol. RNA quality was assessed with Agilent 2100 Bioanalyzer System. Only samples with RNA integrity number (RIN) values between 9.9 and 10 were processed for sequencing. For the RNA-Seq ultra-low input poly-A selection, each sample was processed to generate a dataset (minimum of 8 M reads per sample) at a 50 nt read length in single-end format (1×50) and sequenced in NextSeq 2000 System (Illumina) at the Center for Genomic Regulation (CRG) Core Facility.

Sequence quality was determined with FastQC v0.11.8 software^66^, revealing over 32 million mean reads with median and mean base quality over 32 using Phred+33 quality score. Two reference genome versions were created based on Ensembl *Mus musculus* (GRCm39; https://www.ncbi.nlm.nih.gov/assembly/GCF_000001635.27): the Y chromosome hard masked (Y-masked) and the Y chromosome with PARs hard masked (YPARs-masked) based on Olney *et al.*^67^. Reads were aligned with Hisat2 v2.1.0 ^68^ to the masked versions of the reference genome. Aligned reads were further processed using Samtools v1.21 ^69^ and quantified to gene level using htseq-count v0.11.2 ^70^ and GTF annotation file. Whole-genome alignments were visualized with Integrative Genomics Viewer (IGV)^71^. RNA-Seq data sets can be accessed at the European Nucleotide Archive (ENA) public repository under the following accession numbers (PRJEB56430/ERA18509105).

Differential expression analysis was performed using the Bioconductor package Deseq2 v1.14.0^72^. Differential expressed genes (DEGs) data were processed with custom R scripts (R v4.4) considering genes with q-value < 0.05 and |log2 fold change| > 1 as significantly up- or downregulated. Gene Set Enrichment Analysis (GSEA)^73^ was performed using R packages clusterProfiler^74^ and enrichplot (DOI: <u>10.18129/B9.bioc.enrichplot</u>) for the statistical analysis and visualization of functional profiles of genes and gene clusters. All graphs were plotted with ggplot2 and GOplot R package^75^.

### Statistical Analyses

The researchers were blinded to the sex and genotype at the time of analysis. The number of quantified animals is indicated in each plot; a minimum of three mice were included. The normality of the distribution was tested using the Shapiro-Wilk test in all cases. Each individual test is indicated in the figure legend of the corresponding experiments. All statistical analyses were conducted using GraphPad Prism 9.

The regression analysis was done using the statistical software Stata (StatCorp. 2025. *Stata Statistical Software*: Release 19. College Station, TX:StataCorp LLC), which explains cell counts using as explanatory variables: a categorical variable (baseline variations), the treatment condition (mock or sh*Dmrt2*), and their interaction. This specification is analogous to a variance decomposition.

Statistically significant differences were considered and represented by p-value: *p < 0.05, **p < 0.01 and *** p < 0.001.

---

**Suppl. Figure 1 (related to Figure 1).**
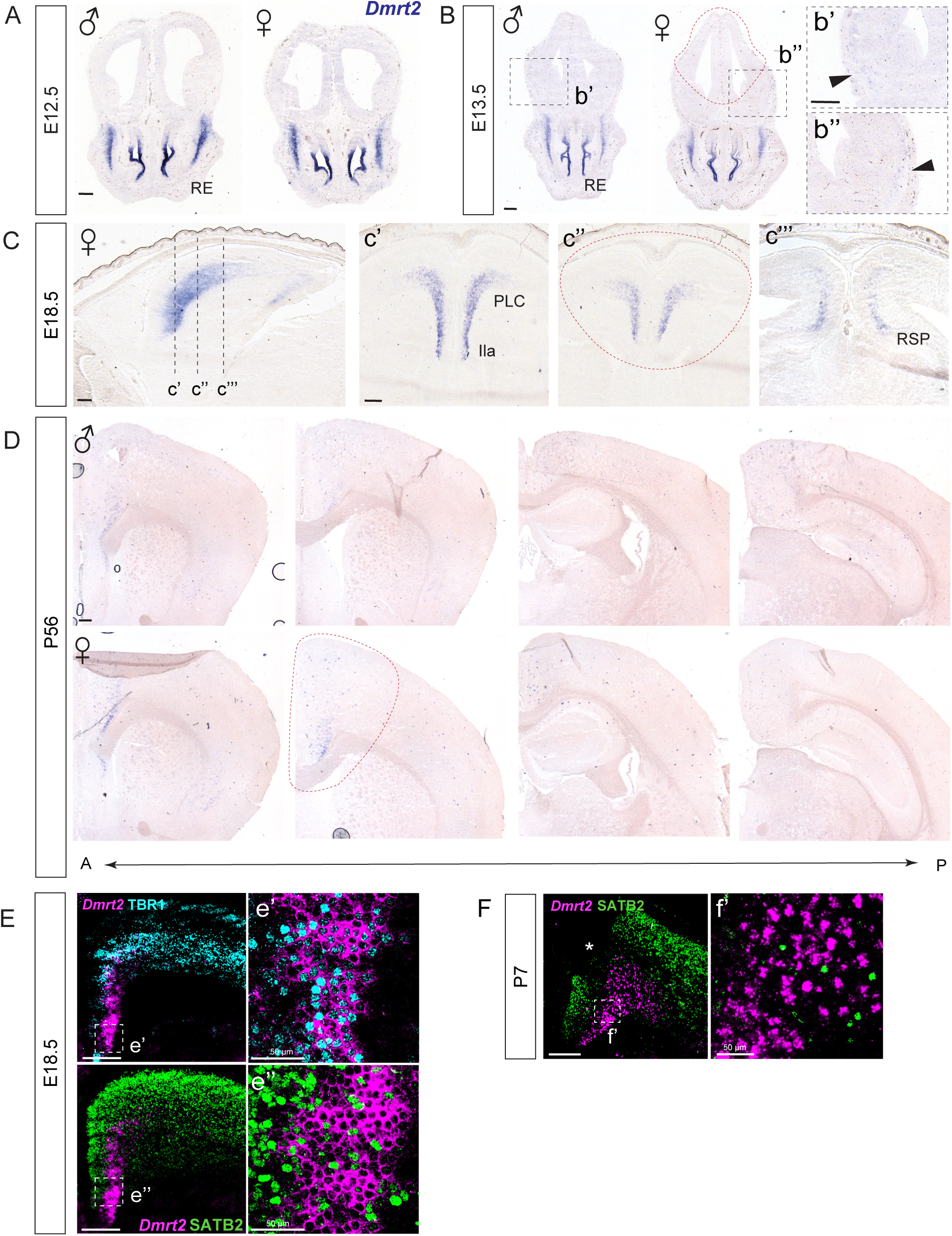
*Dmrt2* embryonic expression in deeper-layer neurons of the mouse cingulate cortex throughout development. ***Dmrt2* ISH in A) male and female E12.5 embryos.** *Dmrt2* expression is clearly detected in the respiratory epithelium (RE). **B) Male and female embryos at E13.5.** A few neurons are detected in the ventral cortex (b’, b’’). Dashed lines indicate the region dissected for qPCR, which includes the cingulate primordium and other regions of the dorsal pallium but excludes the ventral pallium and subpallial regions. **C) Female embryos at E18.5 shows a similar pattern to male embryos**. (c’-c’’’) Coronal sections corresponding to the rostrocaudal positions are depicted in the sagittal view (left panel). **D) *Dmrt2* is maintained in male and female adult animals** along the anteroposterior axis, as coronal views show. A: anterior, P: posterior. **E-F) *Dmrt2* mRNA co-staining with TBR1 and SABT2 at E18.5 (E) and P7 (F).** (e’-e’’, f’) Insets of deep layers of the cingulate cortex. Red lines indicate the dissected area for the qPCR and RT-PCR along the whole rostrocaudal axis. Scale bar: 250 μm, except in the indicated cases. (*) indicates broken tissue.

**Suppl. Figure 2 (related to Figure 1).**
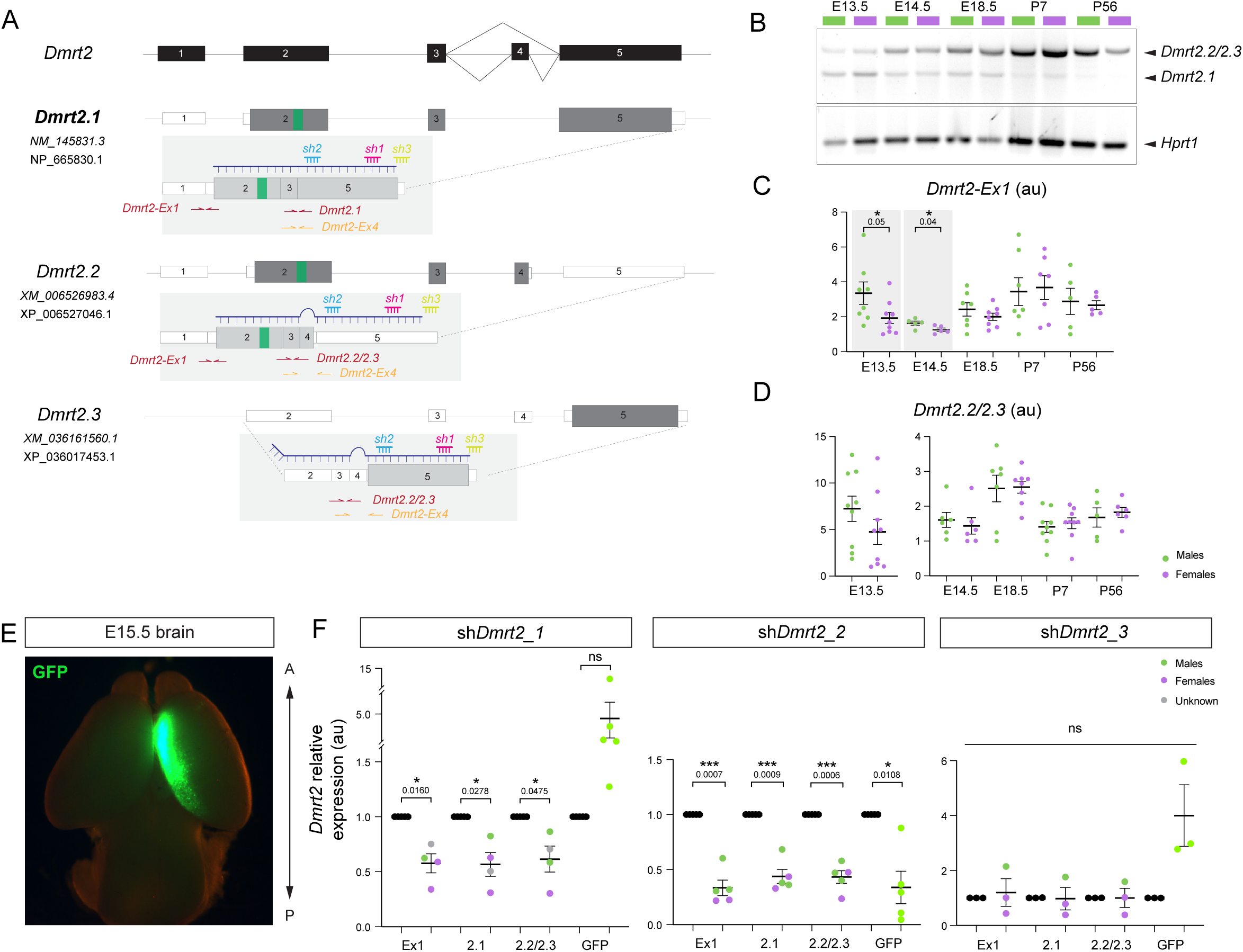
*Dmrt2* splicing variants display a dynamic expression during development. **A) *Dmrt2* locus and the three putative splicing variants** according to NCBI genome browser (Gene ID: 226049). Solid black and grey boxes represent exons; white boxes represent untranslated regions (UTR); green box within exon 2, the DM domain; purple line, *Dmrt2* riboprobe used for ISH detection; red, magenta, and orange arrows represent qPCR primers; blue, yellow, and pink lines represent interference RNAs used in this study to target *Dmrt2.* **B) *Dmrt2* variants switch throughout development.** RT-PCR for *Dmrt2.1* and *Dmrt2.2/2.3* variants across development in male and female. In early embryos, *Dmrt2.1* is the most abundant variant. From E14.5 onwards, *Dmrt2.1* levels are gradually reduced while *Dmrt2.2/2.3* increase. At least three animals were included per time point. **C) *Dmrt2-Ex1*** primers detect variants *Dmrt2.1* and *Dmrt2.2*, but not *Dmrt2.3*. **D) *Dmrt2.2/2.3* relative expression analysis across development.** RT-qPCR with *Dmrt2_Ex1* showed a similar trend to *Dmrt2.1* (Figure 1). There are 1.74 times more *Dmrt2* levels in males than females at E13.5, and 1.30 times more at E14.5. However, no differences were observed between the sexes for *Dmrt2.2/2.3*. Cingulate cortices of males and females were analyzed by qPCR at the indicated time points. Results are the mean±SEM. Each dot corresponds to one individual embryo. Two-tailed t-test: not significant (n.s) and p-value > 0.05 (n ≥ 5); au, arbitrary units. **E) GFP cingulate cortical region in an E15.5 electroporated brain. F) *Dmrt2* reduction after in utero electroporation** with three individual shRNAs. Total *Dmrt2-Ex1*, *Dmrt2.1*, and *Dmrt2.2/2.3,* and *GFP* expression levels were analyzed 5 days after IUE (at E13.5). The contralateral CgCx was used as reference. Each dot corresponds to the ipsilateral (electroporated) side normalized to its contralateral side. Purple dots are female embryos, green dots are male embryos, and grey dots are undetermined. Only sh*Dmrt2*_1 and sh*Dmrt2*_2 showed significant reductions in *Dmrt2* expression. All *Dmrt2* variants were similarly affected by the shRNAs.

**Suppl. Figure 3 (related to Figure 2 and 3).**
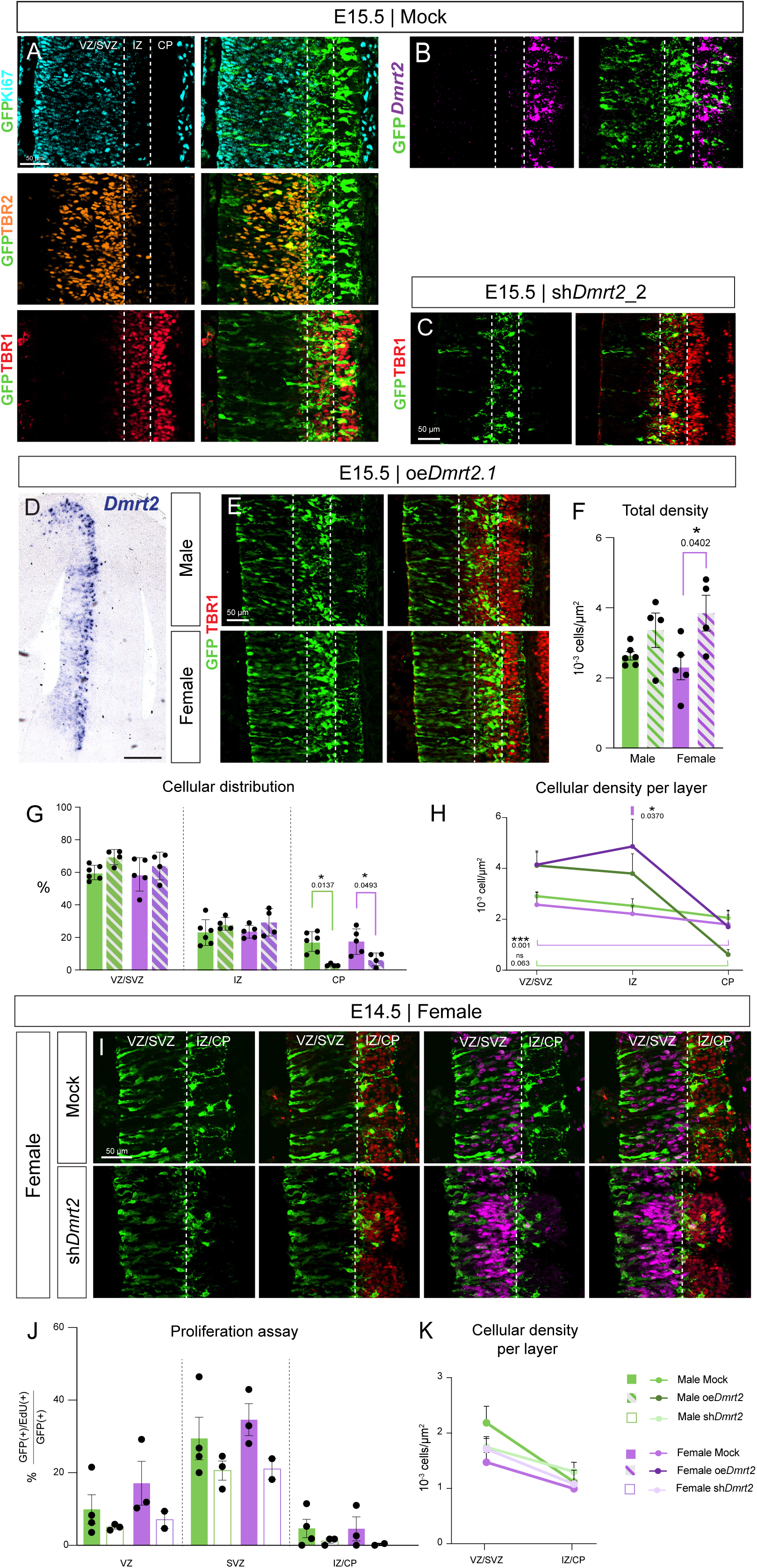
**A) GFP(+) distribution in cingulate primordium layers at E15.5 in female mock**. **B) At E15.5, *Dmrt2*, detected by FISH, is located in the CP**. **C)** GFP distribution in an E15.5 embryo electroporated at E13.5 with a different sh*Dmrt2* (sh*Dmrt2*_2; see Figure 1E for approximate location in *Dmrt2* locus). Similar phenotypes between the two sh*Dmrt2* iRNAs are observed. **D-H) *Dmrt2.1* overexpression (oe) produces the opposite effect to *Dmrt2* downregulation. D) *Dmrt2* ISH in *oeDmrt2.1* E15.5 embryo.** The left hemisphere shows *Dmrt2* signal compared to the non-electroporated contralateral side. The ISH was developed for a short time, such as the endogenous *Dmrt2* is not showing, and only the overexpressed signal is detected. **E) GFP distribution in oe*Dmrt2* male and female CgCx primordia at E15.5. F) Total GFP cellular density within the three layers.** Tukey’s multiple comparison test, ordinary one-way ANOVA (n ≥ 4). **G) Cellular density across each layer in the four conditions.** Dunn’s multiple comparison test, Kruskal-Wallis test (n ≥ 4) **H) Quantifications of GFP(+) cellular distribution (%).** Tukey’s multiple comparison test, ordinary one-way ANOVA (≥ 4). Regression analysis was conducted to compare the cellular density progression between VZ and IZ layers, and VZ and CP layers. **I-K) *Dmrt2* downregulation reduces the percentage of progenitors in both sexes. I) GFP distribution at E14.5 in female embryos. J) Proliferation assay in male and female.** The percentage of GFP+ and EdU+ cells is quantified for mock and sh*Dmrt2* embryos. **K) Cellular density in VZ/SVZ and IZ/CP**. Each dot corresponds to one embryo for all graphs; results are expressed as mean ± SEM and significance mean * p-value < 0.05 and *** p-value < 0.001.

**Suppl. Figure 4 (related to Figure 4).**
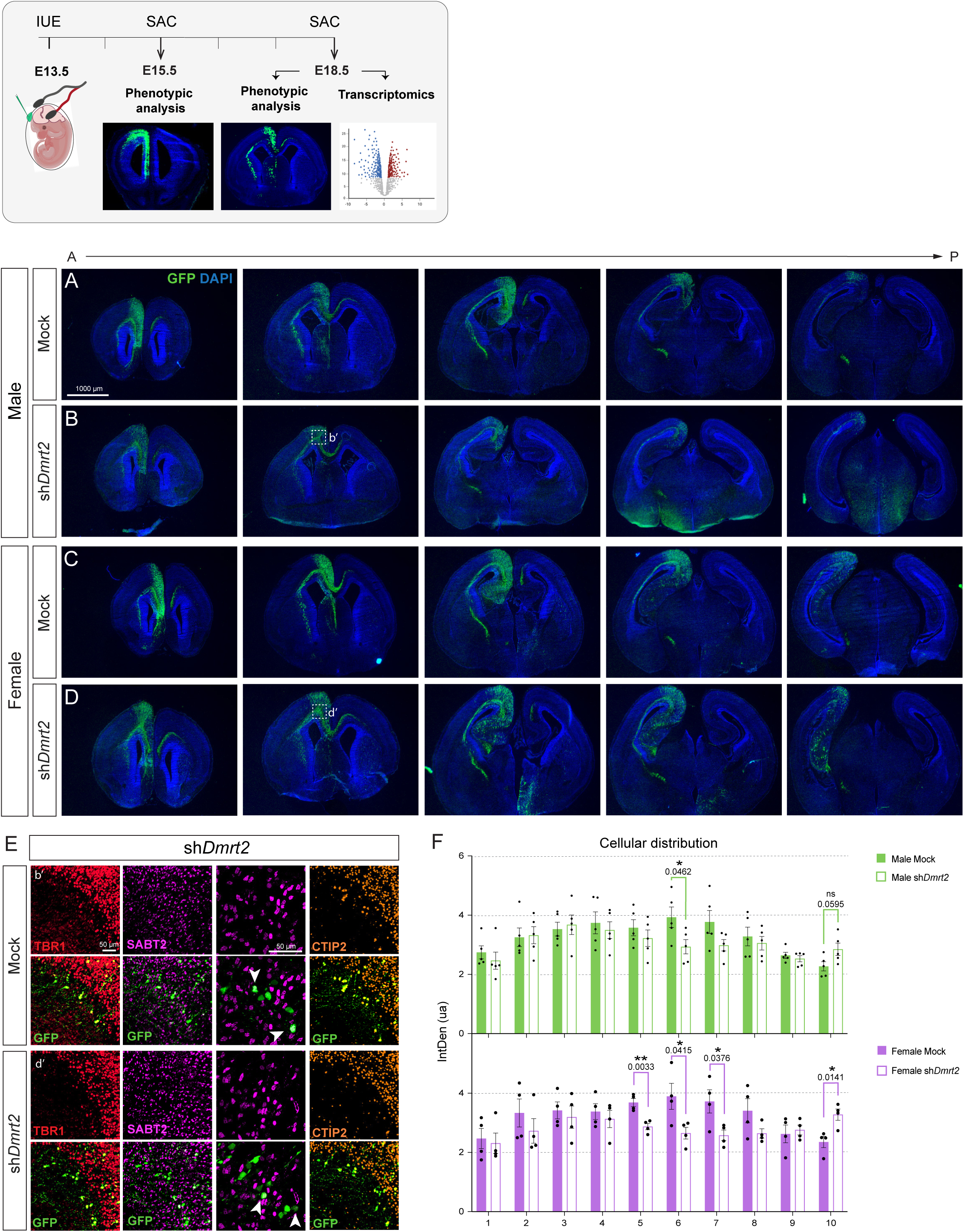
*Dmrt2* silencing causes phenotypic alterations in the CgCx. **Summary of experimental design.** In this study, all brains were electroporated at E13.5 and then analyzed either at E15.5 or E18.5, depending on the experiment. **A-D) Representative sequential sections across the rostrocaudal axis from E18.5 brains, electroporated at E13.5. A) Male mock, B) Male sh*Dmrt2*, C) Female mock, and D) Female sh*Dmrt2*** (A: anterior and P: posterior). **E) GFP(+) cells stacked at the corpus callosum**. All colocalize with TBR1 and CTIP2, but only a few with SATB2. White arrowheads correspond to GFP and SATB2 colocalization. **F) Cellular distribution of GFP(+) cells in the medial CgCx.** Two-tailed t-test (n = 5). Each dot represents an individual embryo. The graph shows the mean±SEM of the average of three sections per embryo, and the significance is as follows: *p-value < 0.05 and **p-value < 0.01.

**Suppl. Figure 5 (related to Figure 5).**
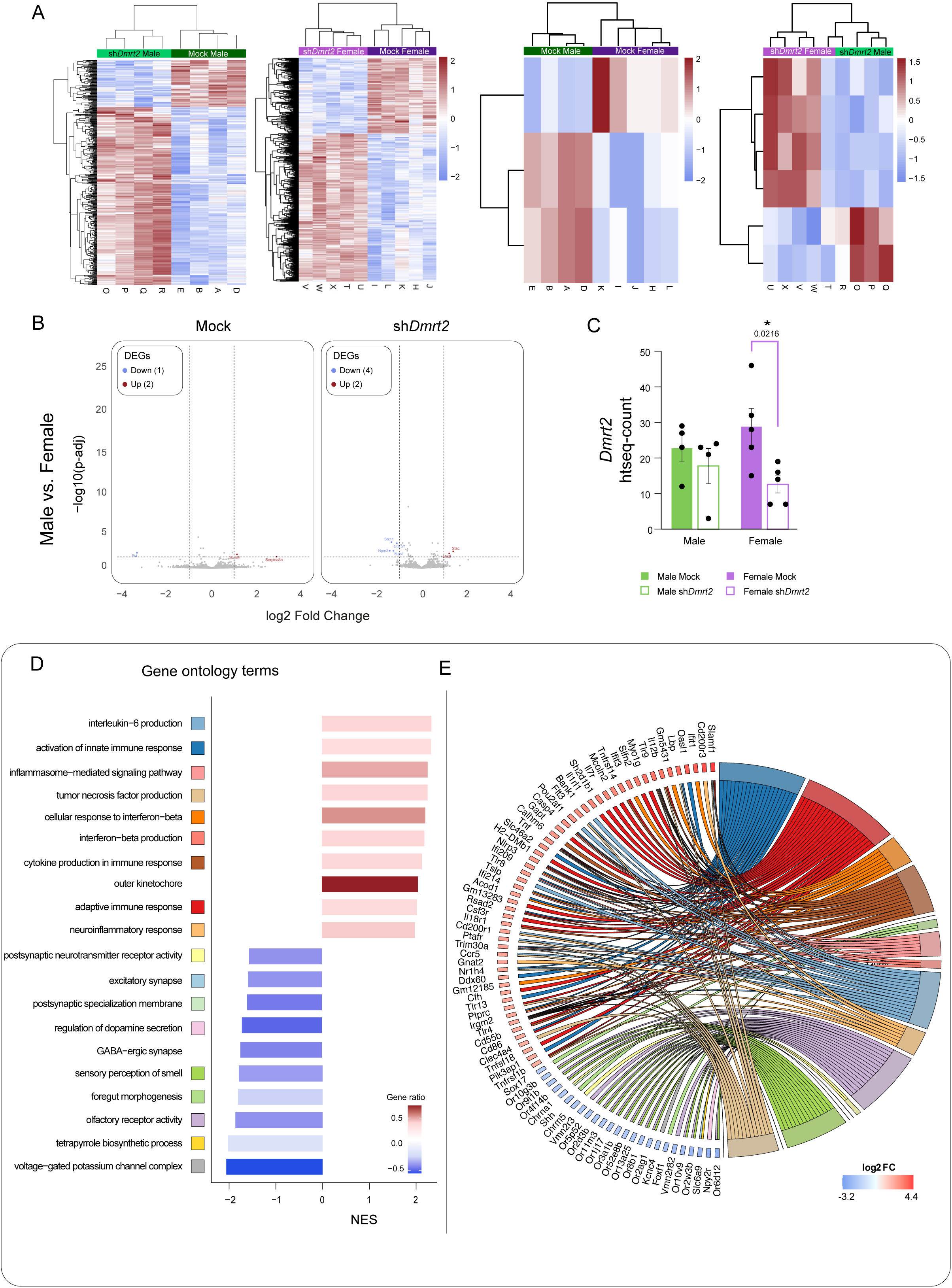
**A)** Heatmaps of DEGs for all comparisons. The color and intensity in the heatmap represent changes in gene expression from the list of genes with |log2FC|<0.5 p-adj < 0.05. **B)** Volcano plots represent DEGs in male vs. female comparisons. Blue color indicates downregulated genes, and red indicates upregulated genes (numbers in brackets). **C)** *Dmrt2* downregulation in RNA-Seq samples. Number of *Dmrt2* reads in the four groups. Two-tailed t-test: * p-value < 0.05 (n ≥ 4). **D-E) Male-specific sh*Dmrt2* vs. mock gene analysis. D)** GSEA for all down- and up-regulated genes. Selected non-redundant categories with a NES > 2 and < 1.5 are shown. **E)** Among the categories depicted in D), genes with the highest |log2FC| between −3.2 and 4.4 are shown in a GO-Chord plot.

## ACKNOWLEDGMENTS

We thank Takahiko Sato for providing the *Dmrt2* probe and Marta Nieto for the *pCAG-GFP* and *pCAGGs-DsRedExpress* plasmids. Prof. Joaquín Artés for statistical regression analysis. The Advanced Light Microscopy, Flow Cytometry, Animal facilities, and the Genomics and NGS Core Facility (GENGS) at the CBM (CSIC-UAM) performed next-generation sequencing data analysis. We also thank Prof. Paola Bovolenta, Dr. Marta Nieto, and lab members, Dr. Jorge García Marqués, and Dr. Leo Beccari for critical reading and advice.

## STATEMENTS & DECLARATIONS

Ethics approval and consent to participate are not applicable.

Written informed consent for publication was obtained from all participants included in the study.

This published article and its supplementary information files include all data generated or analyzed during this study. RNA-Seq raw data are available in the European Nucleotide Archive repository, https://www.ebi.ac.uk/ena/browser/view/PRJEB56430.

Grant PGC2018-101751-A-100 and PID2021-127235NB-I00 funded by MCIN/AEI/ 10.13039/501100011033, and ERDF, “A way of making Europe”.

AB-S holds an FPI fellowship from the Spanish MCIN, PRE19-089366. RTC holds an FPI from the CAM, PIPF-2022/SAL-GL-24909, cofounded by ESF. RC-N holds an FPU from Spanish MCIN, FPU19/02352. ES-S. was supported by Grant RYC-2016-20537 funded by MCIN/AEI/10.13039/501100011033 and ESF, “Investing in your future”. Financial interests: All authors declare they have no financial interests.

## Authors’ contributions and information

ES-S conceived and designed the study and supervised the project. ES-S and AB-S developed the theoretical framework. AB-S performed all the experiments. MR-G contributed to RT-qPCRs, RTC to overexpression analysis, and RC-N to *in situ* hybridizations. All authors discussed the results, ES-S, AB-S, and MR-G contributed to the writing of the manuscript. All authors approved the final version.

